# Leukaemia cell intrinsic and extrinsic factors cooperate to facilitate the survival and proliferation of KMT2A-rearranged B-ALL in the CNS niche

**DOI:** 10.1101/2024.12.03.626602

**Authors:** Alasdair Duguid, Camille Malouf, Leslie Nitsche, Neil A. Barrett, Owen P. Smith, Chris Halsey, Katrin Ottersbach

## Abstract

Infant B-cell acute lymphoblastic leukaemia (B-ALL) is a rare, aggressive entity characterised by KMT2A rearrangements and poor outcomes. One of the unique features of KMT2A-rearranged infant ALL that contributes to these poor outcomes is a particularly high rate of central nervous system (CNS) involvement. There is a broad lack of understanding regarding the molecular processes and immune environment in CNS ALL. This hinders the development of targeted therapies for CNS ALL, which is increasingly essential given the potentially reduced efficacy of current immunotherapies in this specialised environment. In this study, we used an immune-competent KMT2A-AFF1+ infant B-ALL murine model to explore the cell-intrinsic and cell-extrinsic mechanisms that underpin this clinically significant complication. We show novel functional impacts on leukaemia propagating cells following exposure to the CNS niche which results in unique leukaemia repopulation dynamics. Transcriptomic and immune cell profiling by niche showed differences in the immune microenvironment between the CNS and bone marrow niches. The CNS niche demonstrates supressed T cell and macrophage activity which may be part of a wider CNS niche-specific immune escape mechanism. Our transcriptomic comparisons by niche also led us to explore the role of PI3K pathway activation in the propagation of leukaemia cells within the CNS niche. We identify miR-93 as a possible master regulator of this process. The importance of miR-93 is emphasised as it is shown to be upregulated in CNS leukaemia cells across multiple murine KMT2A-AFF1 B-ALL model systems and in primary KMT2A-AFF1 B-ALL patient samples. We conclude by showing impaired CNS engraftment of leukaemia cells upon miR-93 knockdown, cementing its importance in CNS leukaemia biology and opening up new CNS-specific therapeutic opportunities.

## Introduction

Acute lymphoblastic leukaemia (ALL) involvement of the central nervous system (CNS) is the most common and most clinically significant extra-medullary site of disease across all age groups. However, there is a patient group at a particularly high risk of this complication: those with KMT2A-AFF1+ infant B-ALL. Compared with non-infant children, the rates of detectable CNS involvement are 8-10 fold higher for infant ALL patients at diagnosis and relapse despite universally adopted CNS-directed therapy throughout upfront treatment(Pieters et al. 2007, 2019; Thastrup et al. 2022). There is no standardised treatment for the management of CNS relapses other than a combination of multi-agent chemotherapy, intrathecal therapy and cellular therapy, including haematopoietic stem cell transplant. For infant patients, survival following the first relapse is very poor at around 20%, and for survivors, CNS-directed therapy is associated with long-term neurocognitive disabilities(Driessen et al. 2016; Iyer et al. 2015). There is a lack of understanding of the biological mechanisms that account for the high rate of CNS involvement in infant ALL. Overcoming this knowledge gap is necessary as part of a broader and urgently needed effort to develop more effective, less toxic therapies to improve the outcomes for these patients.

Experiments using primary childhood B-ALL patient samples and patient-derived xenograft models have shown that it is a generic ability of leukaemia cells to enter the leptomeninges (the site of CNS disease in ALL)(Williams et al. 2016; Bartram et al. 2018). Once in the CNS niche, they must adapt their metabolism to the different environmental conditions, most notably low fatty acid levels, low cholesterol and relative hypoxia (Savino et al. 2020; Cousins et al. 2022; Kato et al. 2017), This study aimed to identify novel transcriptomic and functional differences between CNS and bone marrow (BM) leukaemia cells unique to KMT2A-AFF1+ infant B-ALL. We speculate that these insights may provide a biological explanation for the clinical finding of high levels of CNS involvement experienced by this patient population and identify new CNS-specific therapeutic targets. We undertook this work primarily using a previously published fully murine model of Kmt2a-AFF1+ infant B-ALL in which overexpression of miR-128a in Kmt2a-AFF1+ fetal liver (FL) Lineage^-^Sca1^+^c-Kit^+^ (LSK) generated an aggressive pro-B ALL phenotype on serial transplantation through immunocompetent mice, (Malouf et al. 2021). Critical for this study, the leukaemia phenotype included prominent CNS involvement in a leptomeningeal distribution mirroring the human disease.

The CNS immune niche in B-ALL remains poorly characterised, a knowledge gap that has become increasingly relevant with the growing reliance on immunotherapies in infant B-ALL treatment, which have largely unknown CNS activity. Our study identified CNS-specific differences in leukaemia cell surface proteins involved in interactions with immune cells, suggesting altered immune dynamics in this niche. Further analysis revealed differences in macrophage and T-cell populations between the CNS and BM leukaemia niches, suggesting reduced anti-leukaemia immune activity within the CNS. There are significant bidirectional interactions between the tumour immune microenvironment and PI3K/Akt in tumour cells(Mafi et al. 2021). We found this pathway to be specifically activated in CNS leukaemia cells. This activation was linked to reduced expression of PTEN and CDKN1A in CNS leukaemia cells and potentially mediated by the upregulation of their direct inhibitor, miR-93. This upregulation was consistently observed across multiple model systems and patient-derived samples.

Importantly, knockdown of miR-93 selectively impaired CNS leukaemia engraftment, highlighting its critical role in CNS-specific leukaemia survival. These data highlight novel CNS niche-specific leukaemia immune regulation and growth factor signalling pathways, which we believe pave the way for targeted CNS-specific interventions for infant B-ALL.

## Results

### Exposure to the CNS niche alters the repopulation phenotype of leukaemia

The BM LPCs were established in the initial publication describing the miR-128a Kmt2a-AFF1+ model, where LSK_IL7R and LK/CLP propagated the disease through serial transplantation(Malouf et al. 2021). However, the function of these LPCs when derived from the CNS niche is currently unknown. Our aim was, therefore, to identify whether CNS-derived LPCs have the same propagating capacity to cause systemic disease as BM-derived LPCs. This will provide a basis to understand whether CNS-derived LPCs are able to drive a systemic relapse, a previously unexplored but potentially treatment altering leukaemia relapse dynamic.

We began by comparing the leukaemia composition in the BM and CNS niche in the miR-128a Kmt2a-AFF1+ model. We transplanted BM-derived LSK_IL7R and LK/CLP separately (Supplemental Table 1) and assessed the resulting leukaemia phenotype in the BM and CNS niches. As shown in Supplemental Figure 1, there were no differences in leukaemia phenotype by type of LPC transplant, and transplant-derived leukaemic cells accounted for >90% of the CD45+ cells in the BM at the terminal leukaemia stage. The terminal leukaemia phenotype was indistinguishable by type of LPC transplanted, and the two different LPCs were able to give rise to each other. Both LPCs gave rise to CNS involvement in a leptomeningeal distribution without difference by LPC type transplanted, example histology images are shown in Figure 1A. The composition of the leukaemia infiltrated in the CNS niche mirrored that in the BM niche (Figure 1B).

**Figure 1.**
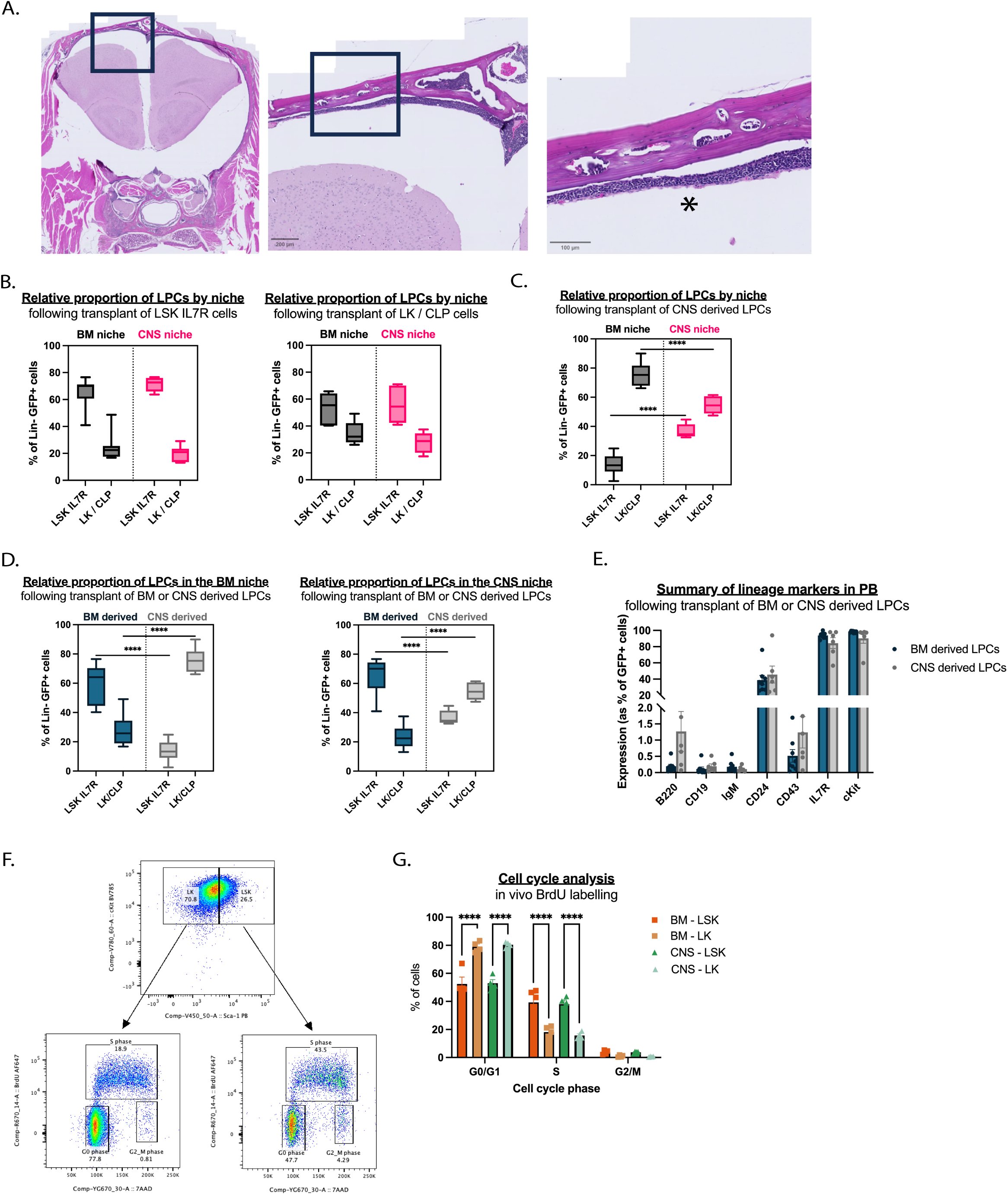
Exposure to the CNS niche alters the repopulation phenotype of leukaemia. (A) Illustrative image of a coronal histology section of the head from a mouse that has developed terminal leukaemia following transplant with BM-derived LSK_IL7R cells. The (*) is adjacent to a dense leptomeningeal leukaemia infiltrate. (B) Comparison of relative proportions of LSK_IL7R and LK/CLP in BM and CNS by type of LPC transplanted (BM-derived LSK_IL7R or LK/CLP). There were no significant differences by type of LPC transplanted in LPC repopulation. Both transplanted cell types repopulated equal proportions of LPCs when comparing the BM and CNS niches. LSK_IL7R transplants N=7, LK/CLP transplants N=5. (C) Comparison of LSK_IL7R and LK/CLP LPCs by niche in CNS-derived LPC transplants. There was an increase in LSK_IL7R and a reduction in LK/CLP in the CNS niche compared with the BM niche. Differences in the relative proportions of LSK_IL7R and LK/CLP in either niche were significant to p<0.0001, BM niche N=8, CNS niche N=5. (D) Comparison of LPCs in BM and CNS niche by the source of the transplanted cells, showing differences in the relative proportions of LPCs in either niche by source of transplanted cells to a significance of p<0.0001. BM niche; BM-derived transplants N=12, CNS-derived transplants N= 8. CNS niche; BM-derived transplants N=12, CNS-derived transplants N= 5. (E) No differences in comparisons of lymphoid marker expression in PB of mice who developed terminal leukaemia by the source of transplanted cells. BM-derived transplants N=8, CNS-derived transplants N= 6. (C-E) Data from LSK_IL7R and LK/CLP cell transplants were pooled for these comparisons, comparing initial transplants from BM-derived LPCs with CNS-derived LPCs (next round recipients). (F) Illustrative cell cycle phase analysis example: prior gating on single GFP+ cells. LSK_IL7 and LK/CLP equivalents GFP+cKit+Sca1+ and GFP+cKit+Sca1-. (G) Comparison of LPCs in both BM and CNS niche by cell cycle phase. A higher percentage of LSK_IL7R is in the S phase, and a lower percentage in the G1 phase in both niches than LK/CLP. N=4 per LPC in each niche. p < 0.0001 for comparisons shown.

We next assessed whether LPCs recovered from the CNS could re-engraft systemically to model the dynamics of a CNS-to-BM relapse event. Both populations of CNS-derived LPCs (LSK_IL7R and LK/CLP) systemically engrafted and resulted in terminal leukaemia. Overall assessment of disease latency, spleen and liver weights, and peripheral blood (PB) counts did not vary by source (BM or CNS) of LPC transplanted (Supplemental Figure 2). However, upon more detailed comparisons between the initial transplants from BM-derived LPCs with CNS-derived LPCs from the subsequent transplant (Supplemental Figure 3), two significant differences were identified. First, in transplants of CNS-derived cells, unlike transplants of BM-derived cells, the composition of the resulting leukaemia differed in the BM and CNS niches by relative proportions of LPCs (Figure 1C). Second, the relative proportions of the two LPCs varied in both niches by the source of transplanted cells upon terminal leukaemia development; transplants of BM-derived cells reconstituted an LSK_IL7R predominant leukaemia phenotype in the initial transplants, whereas CNS-derived cells reconstituted an LK/CLP predominant leukaemia phenotype when re-transplanted systemically (Figure 1D). There was no difference in lineage-positive PB output by the source of transplanted LPC (Figure 1E). These data suggest that LPCs experience lasting changes after exposure to the CNS environment. These functional changes become apparent on relocation to and expansion in the BM niche.

We speculated that the differences in repopulation by the source of LPC might be due to inherent cell cycling differences between LPCs and a bias in the CNS niche towards enriching for a quiescent phenotype(Jonart et al. 2020). Using *in vivo* BrdU labelling and additional cell surface staining, we assessed the cell cycling status of LPCs from the BM and CNS niche. Figure 1F-G shows that cKit+Sca1+ cells (equivalent to LSK_IL7R cells) are more actively cycling than cKit+Sca1-cells (equivalent to LK/CLP cells) in the BM and CNS niches. Although comparisons between niches may be limited due to differences in BrdU distribution, there were no apparent differences in cell cycle by niche.

### Transcriptomic analyses reveal cycling and metabolic adaptations to the CNS niche

To identify differences between BM and CNS LPCs on a transcriptional level, we generated bulk RNA sequencing data of paired BM and CNS LPCs. As an initial validation, we performed comprehensive transcriptomic comparisons against several published datasets. Comparing the differentially expressed genes (DEGs) of BM-derived LPCs from our model to those from a dataset that consists of 21 patients with KMT2A-AFF1+ B-ALL, 16 infant and five non-infant paediatric cases(Andersson et al. 2015), the gene expression of the murine model clearly clusters with infant patient samples. This establishes our model as representative specifically of infant disease rather than paediatric KMT2A-AFF1+ B-ALL (Supplemental Figure 3A). Furthermore, its fetal origin was confirmed through comparison against our published RNA sequencing data from wild-type murine FL and BM lymphoid-primed multipotent progenitors (LMPPs)(Symeonidou et al. 2021). This showed that the gene expression of both LPC (LK/CLP and LSK_IL7R) populations from the miR-128a Kmt2a-AFF1+ model was more similar to FL LMPPs rather than BM LMPPs (Supplemental Figure 3B). FL LMPP and miR- 128a Kmt2a-AFF1+ samples shared gene expression patterns of many established onco-fetal genes in infant leukaemia, namely Lin28b, Igf2bp1/2/3 and Hmga2 (Supplemental Figure 3C)(Busch et al. 2016; He et al. 2018; Palanichamy et al. 2016; Roy et al. 2013; Symeonidou et al. 2021).

Further analyses between LPC populations and niches in our newly generated dataset, shows 52 DEGs between BM LSK_IL7R and LP/CLP cells (Figure 2A) with enrichment for cell cycle-related gene ontologies, including the upregulation of genes associated with proliferation, such as *Kpna2*, in LSK_IL7R cells (Han and Wang 2020). Figure 2B shows that LPCs in the CNS niche were transcriptionally more similar, with fewer DEGs, than in the BM niche. Lists of these DEGs are available in Supplemental Tables 2 and 3. The only commonly differential expressed gene between LPCs in both niches was the cell cycle gene *Ube2c*, which was upregulated in LSK_IL7R cells .

**Figure 2.**
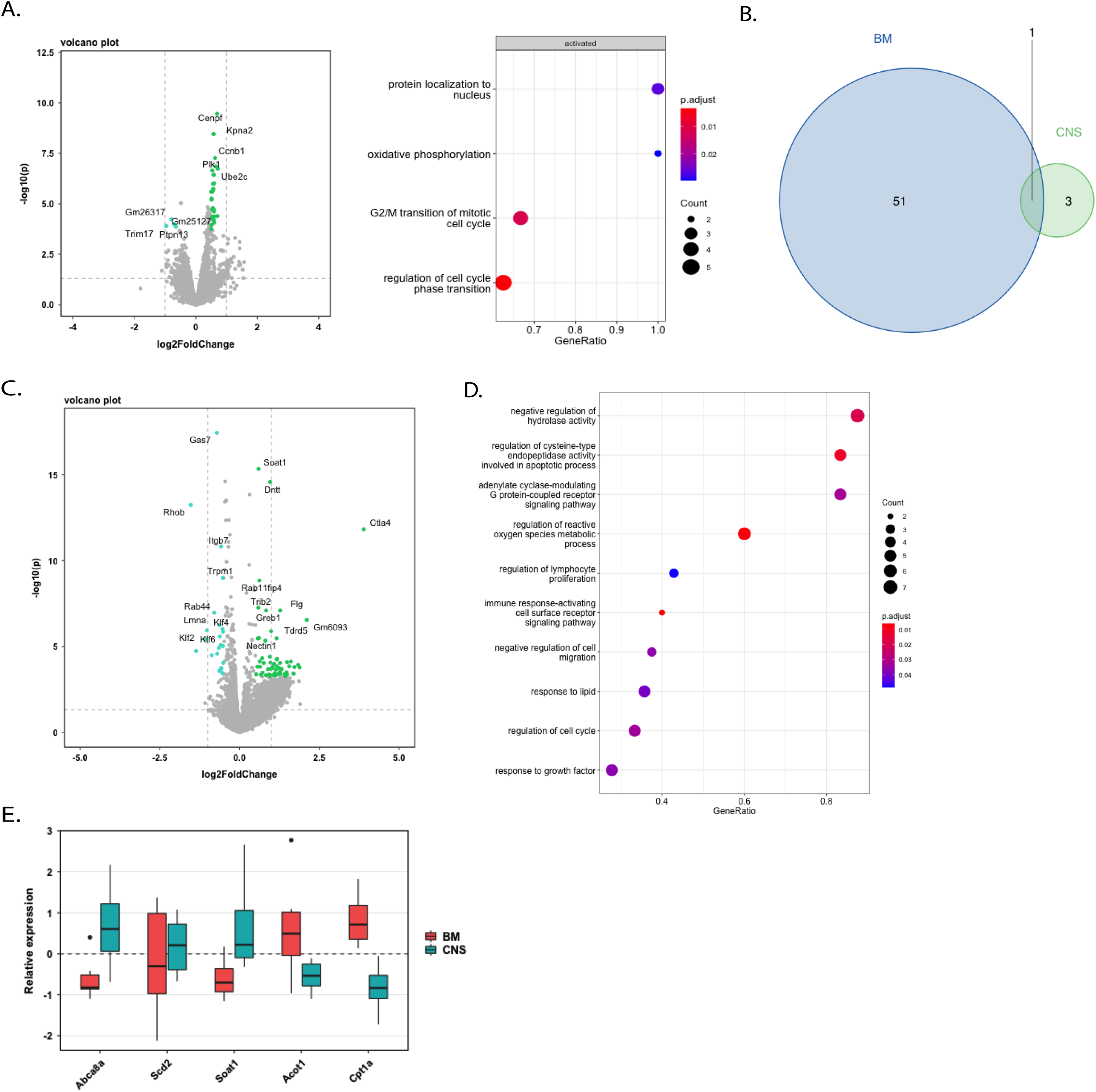
Transcriptomic analyses reveal cycling and metabolic adaptations to the CNS niche. (A) Volcano plot of DEGs between LSK_IL7R and LK/CLP cells from the BM. Genes upregulated in LSK_IL7R in green and downregulated in teal; p.adj<0.05 log2FC threshold 0.5. Enriched gene ontologies, many of which are cell cycle related, in BM LSK_IL7R cells relative to BM LK/CLP cells are also shown. (B) Venn diagram showing the number of DEGs between LPCs in the BM (blue) and CNS (green) niche. Ube2c is the single overlapping DEG. (C) Differentially expressed genes by niche, where genes upregulated (green) and downregulated (teal) in the CNS-derived LPCs were compared with BM-derived LPCs using p.adj <0.05 and log2FC 0.5. The LPCs - LSK_IL7R and LK/CLP - were pooled for this analysis. The top 10 upregulated and downregulated genes by adjusted p-value are labelled. (D) Differentially regulated biological processes by GO terms between CNS and BM-derived LPCs. (C-D) Comparisons made from the analysis of 8 independent paired BM and CNS LPC samples. (E) Scaled normalised read counts of differentially expressed genes involved in lipid metabolism by niche with p.adj < 0.05. Abca8a: ATP Binding Cassette Subfamily A Member 8; Scd2: Stearoyl-CoA Desaturase-2; Soat1: Sterol O-Acyltransferase 1; Acot1: acyl-CoA thioesterases; Cpt1a: Carnitine Palmitoyltransferase 1A.

We extended our comparisons of CNS LPCs in the miR-128a Kmt2a-AFF1+ B-ALL model to include patient CSF microarray data and patient-derived xenograft CNS B-ALL data(Van Der Velden et al. 2016; Savino et al. 2020). Figure 2C-D summarises the gene expression differences between pooled LPCs derived from BM and CNS in the miR- 128a Kmt2a-AFF1+ B-ALL model; a full list of significant DEGs can be found in Supplemental Table 4. Our dataset revealed CNS-specific adaptations to lipid metabolism previously described in paediatric B-ALL, including the upregulation of SCD (murine ortholog Scd2 in our dataset). Additional metabolic adaptations to the CNS niche involving genes associated with fatty acid and cholesterol metabolism found in our model are summarised in Figure 2E. These CNS-leukaemia metabolic adaptations, described here for the first time in infant leukaemia, have been shown to be essential for B-ALL cell survival, growth, and proliferation within the CNS niche, which is characterised by limited availability of environmental fatty acids.

### Niche-specific leukaemia-immune cell interactions and immune cell populations

A group of DEGs between LPCs from the CNS and BM-niche indicate niche-specific regulation of leukaemia-immune cell interactions, including with T cells, NK cells, and macrophages. The most highly upregulated gene when comparing CNS-derived with BM-derived LPCs is *Ctla4* (p.adj <0.001, log2FC 3.89), illustrated in Figure 2C. This immune checkpoint is well characterised when expressed by T cells, where it suppresses T cell function and proliferation. The immune checkpoint and stem cell marker *Cd96* was also found to be upregulated by CNS-derived LPCs (p.adj 0.001, log2FC 0.57). *Cd47* encodes a macrophage immune checkpoint and is downregulated in CNS-derived LPCs (p.adj 0.001, log2FC 0.19).

There are also leukaemia cell-intrinsic differences in the transduction of immune signalling. Several genes involved in B cell receptor (BCR) signalling were differentially expressed by niche: *Cd22*, *Syk* and *Bcl6*. *Cd22*, which was upregulated in CNS-derived LPCs (p.adj 0.048, log2FC 1.43), is expressed on the surface of B cells from the immature B cell stage; however, it is also expressed by pro-B ALL cells (Lanza et al. 2020). CD22 binds α-2,6-linked sialic acid expressed on B and T cells and some endothelial cells and has a role in modulating intracellular signalling from the BCR with a predominantly inhibiting effect. Associated with a reduction in the propagation of BCR signalling, *Syk*, an essential mediator of this signal, was downregulated in CNS-derived cells (p.adj 0.004, log2FC 0.31). *Bcl6*, upregulated in CNS-derived cells (p.adj 0.014, log2FC 0.63), also modulates BCR signalling and promotes B-cell survival.

Based on these DEGs, we performed targeted immunophenotypic profiling by FACS of these immune cells by niche. We characterised the T cell contribution to the niches by subset and assessed for makers of activity. Paired comparisons of T cell subsets from the CNS and BM leukaemia niches found an equal number of total T cells per 100,000 leukaemia cells, but with fewer CD4+ T cells and T regulatory cells and more CD8+ T cells in the CNS niche (Figure 3A). There was an almost 4-fold increase in the proportion of CTLA4+ T cells in the CNS leukaemia niche compared with the BM, implying a more inhibited T cell population (Figure 3B). This was associated with a nearly 5-fold increase in the proportion of PD-1+ CD8+ T cells in the CNS leukaemia niche (Figure 3B). PD-1 is also an immune checkpoint and a T cell exhaustion hallmark (Y. Jiang, Li, and Zhu 2015). Compared with CTLA4, PD-1 inhibits T-cell function at a later stage following activation (Buchbinder and Desai 2016). In addition, there was reduced expression of the early T cell activation marker CD69 by CD4+ cells in the CNS niche (Figure 3B).

**Figure 3.**
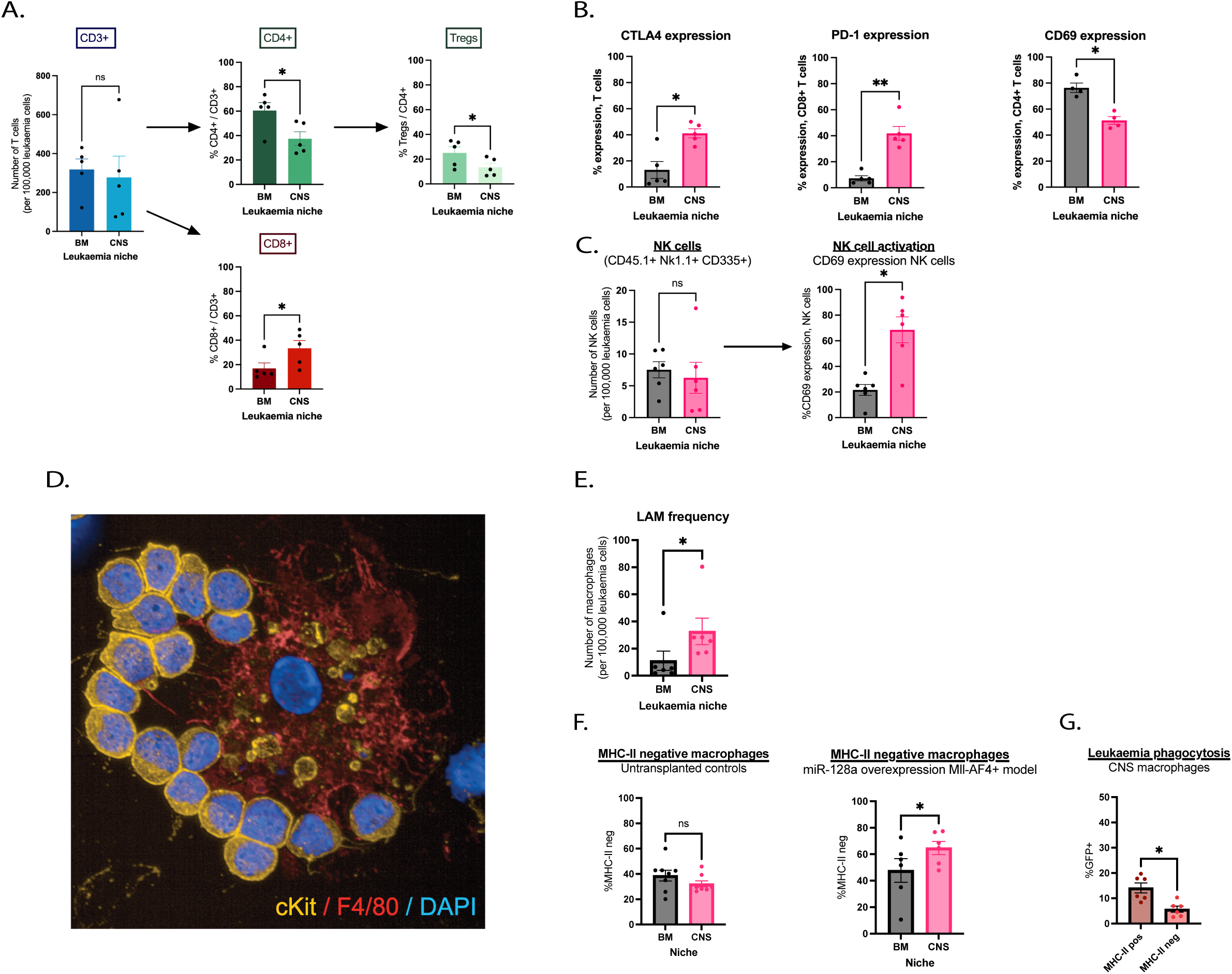
Niche-specific leukaemia-immune cell interactions and immune cell populations. (A) Comparing T cell subsets in paired analyses from the CNS and BM leukaemia niche. Equal frequency of T cells per 100,000 leukaemia cells in either leukaemia niche, p=0.76. CD4+ T cells were lower, and CD8+ T cells were higher in proportion to total T cells in the CNS niche compared with the BM niche, p=0.01 and p=0.03, respectively. As a proportion of CD4+ T cells, T regulatory cells (CD3+ CD4+ CD25+ Foxp3+) were lower in the CNS niche compared with the BM niche, p=0.02. N=5 paired samples. (B) Higher proportion of CTLA4 positive T cells in the CNS than BM niche, N=5 paired samples, p=0.01. Higher proportion of PD-1 positive CD8+ T cells in the CNS than BM niche, N=5 paired samples, p<0.01. Reduced proportion of CD69 positive CD4+ T cells in the CNS than BM leukaemia niche, N=4 paired samples, p<0.02. (C) Equal frequency of NK cells per 100,000 leukaemia cells in either CNS and BM niche. The proportion of CD69+ NK cells was higher in the CNS leukaemia niche, p=0.01. N=6 paired samples. (D) Leukaemia-associated macrophage (LAM) cytospin image of whole CNS infiltrate stained with fluorescently labelled antibodies against cKit (yellow, leukaemia cell) and F4/80 (red, macrophage). (E) LAM (CD45.1+ F4/80+ Ly6C/6G- CD11b+ IA/IE+) frequency per 100,000 leukaemia cells was higher in the CNS than BM niche. N = 6 paired samples, p=0.031. (F) MHC-II negative macrophages predominate in the CNS leukaemia niche. MHC-II negative macrophages were identified as the IA/IE- fraction of CD45.1+ F4/80+ Ly6C/6G- CD11b+ IA/IE+/- cells and expressed as a percentage thereof. In age-matched (8-12 weeks old) untransplanted controls, there was no difference in the percentage of MHC-II negative macrophages by niche, n=8 paired samples, p=0.20. In the miR-128a overexpression model, macrophages were more likely to be MHC-II negative in the CNS niche, n=6 paired samples, p=0.02. (G) Comparison of GFP+ or ’actively phagocytic’ MHC-II positive and negative macrophages. MHC-II negative macrophages are less phagocytic in the CNS leukaemia niche. N=6 paired samples, p=0.01.

These data provide evidence of increased T-cell inhibition at both early and late stages of activation in the CNS leukaemia niche. This pattern of cell surface marker should be associated with immune tolerance and reduced T cell-mediated cytotoxic activity in the CNS leukaemia niche.

We made a similar comparison with NK cells (Figure 3C), which have been shown to have unique properties in the meningeal niche(Garofalo et al. 2023). NK cells (GFP- CD45.1+ CD3/19/Ter119- Nk1.1+ CD335+) in the CNS and BM leukaemia niche were identified by flow cytometry along with CD69 expression which, as with T cells, is an early activation marker on NK cells and a surrogate marker for anti-cancer effector function(Frishman-Levy et al. 2015). There were equal numbers of total NK cells per 100,000 leukaemia cells; however, a higher representation of NK cells in the CNS expressing the activation marker CD69.

Finally, we characterised leukaemia-associated macrophages (LAM). LAMs were easily identified morphologically and by immunofluorescence of cytospin preparations of the CNS and BM infiltrate (Figure 3D), which showed ckit+ leukaemia cells tightly associated with F4/80+ macrophages that showed signs of phagocytosed ckit+ leukaemia cell remnants. Macrophages (defined as GFP- CD45.1+ F4-80+ Ly6C/G- CD11b+ MHC-II+) were more frequent in the CNS than the BM niche per 100,000 leukaemia cells in a paired comparison (Figure 3E). We also focused on an MHC-II- negative macrophage population described in Rebejac et al.(Rebejac et al. 2022). They described that the balance between MHC-II positive and negative macrophages in the meninges was found to change with age, with younger age associated with an MHC-II negative enriched macrophage population, and function, with MHC-II positive macrophages having a more inflammatory anti-microbial phenotype. For this comparison, we defined these populations as CD45.1+ F4/80+ Ly6C/6G- CD11b+ cells and either MHC-II positive or negative. MHC-II negative macrophages were found to account for a larger percentage of total macrophages in the CNS compared with the BM niche in leukaemic mice, but not untransplanted controls (Figure 3F). To assess the function of MHC-II negative macrophages, GFP positivity was used as a surrogate marker for phagocytosis of GFP+ leukaemia cells. MHC-II negative macrophages demonstrated less leukaemia phagocytosis in the CNS niche than their MHC-II positive counterparts (Figure 3G). Based on this evidence, we hypothesise that MHC-II negative macrophages, which become enriched in the meninges in the context of leukaemia, may perform a supportive rather than inflammatory function.

### Kmt2a-AFF1+ infant B-ALL in the CNS niche has increased activation of PI3K pathway signalling relative to leukaemia in the BM niche

Transcriptomic comparisons between LPCs from the BM and CNS demonstrated differential regulation of growth factor signalling (Figure 4A), involving two receptor tyrosine kinases with upregulation of *Met* (p.adj 0.022, log2FC 1.88) and downregulation of *Pdgfrb* (p.adj <0.001, log2FC 0.62) in CNS-derived LPCs, although only c-MET was found to be differentially expressed on a protein level (Figure 4B). One of the pathways activated downstream of these receptors is the PI3K pathway(Engelman, Luo, and Cantley 2006; Hubbard and Miller 2007), with two associated regulators, Pten and Cdkn1a, also being differentially expressed. Pten has an opposing action to PI3K by preventing the formation of PIP3 (Figure 4A), effectively blocking signal transduction, and was downregulated in CNS-derived cells (p.adj = 0.0003, log2FC 0.2). Cdkn1a, which was also downregulated in CNS LPCs (p.adj = 0.004, log2FC 1.4), blocks cell cycle progression and is inhibited by activation of the PI3K pathway through the action of phosphorylated AKT(Freedman, Wu, and Levine 1999; Engeland 2022; Shamloo and Usluer 2019).

**Figure 4.**
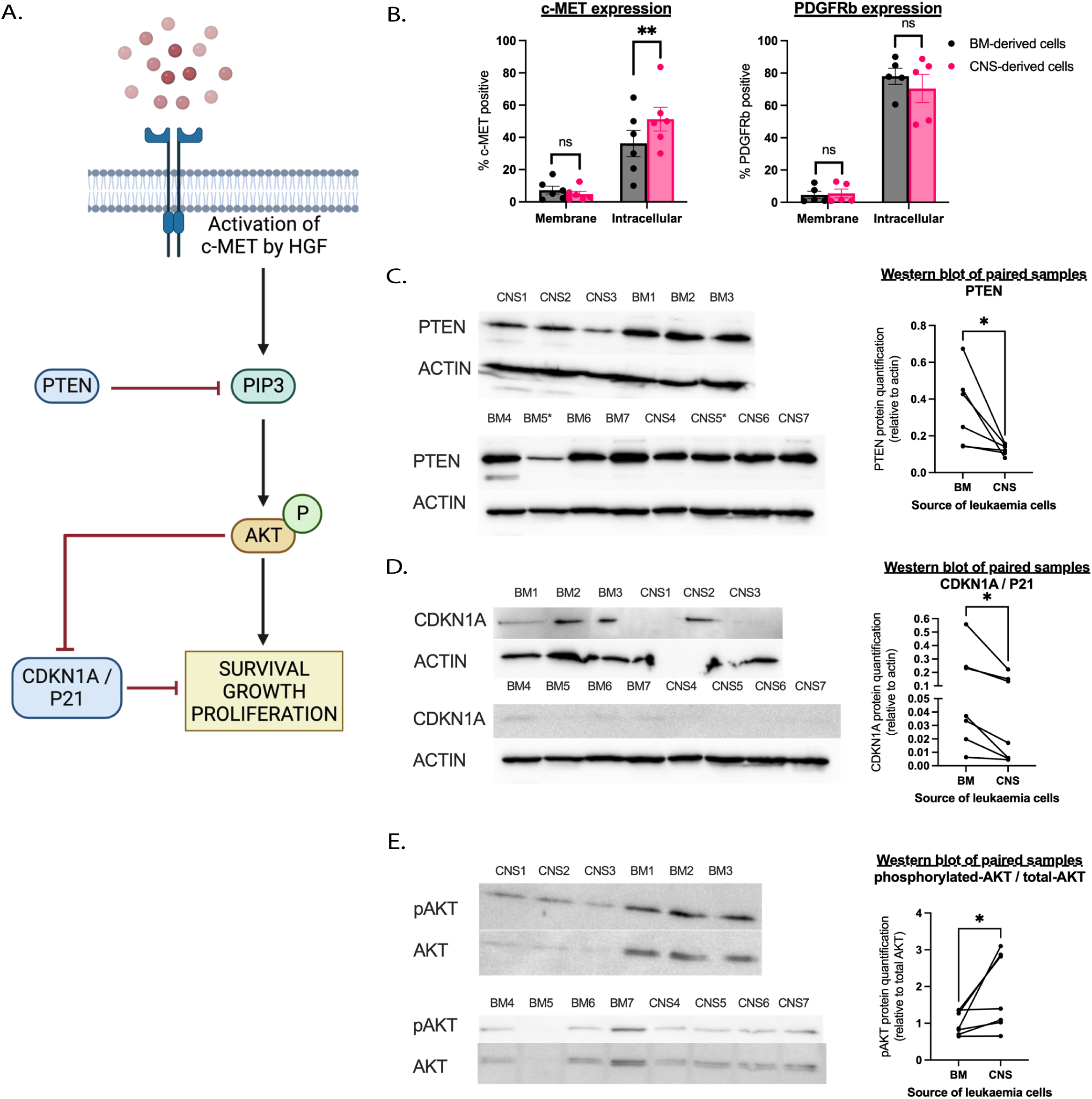
Kmt2a-AFF1+ infant B-ALL in the CNS niche has increased activation of PI3K pathway signalling relative to leukaemia in the BM niche. (A) Summary of PI3K pathway highlighting the roles of c-MET, PTEN and CDKN1A. (B) Flow cytometry quantification of c-MET and PDGFRβ positive LPCs. N=6 paired samples for c-MET comparison, N=5 paired samples for PDGFRβ comparison. p<0.01 for the intracellular c-MET comparison. (C-E) Protein quantification of CNS and BM-derived cells, N=7 paired samples. Western blot images cropped to focus on the protein of interest band and relative quantification are presented. (C) PTEN quantification with N=6 paired samples, p = 0.04. Replicate BM5 and paired sample CNS5 (marked with asterisks(*)) was identified as an outlier and excluded as the quantification was approximately 10- fold lower than the mean. (D) CDKN1A quantification with N=7 paired samples, p = 0.02. (E) Phosphorylated AKT quantification relative to total AKT, N=7 paired samples, p=0.047.

We validated Pten and Cdkn1a downregulation on a protein level in extracts from whole meningeal cell infiltrate compared with whole BM infiltrate in mice with terminal leukaemia (Figures 5C-D).

**Figure 5.**
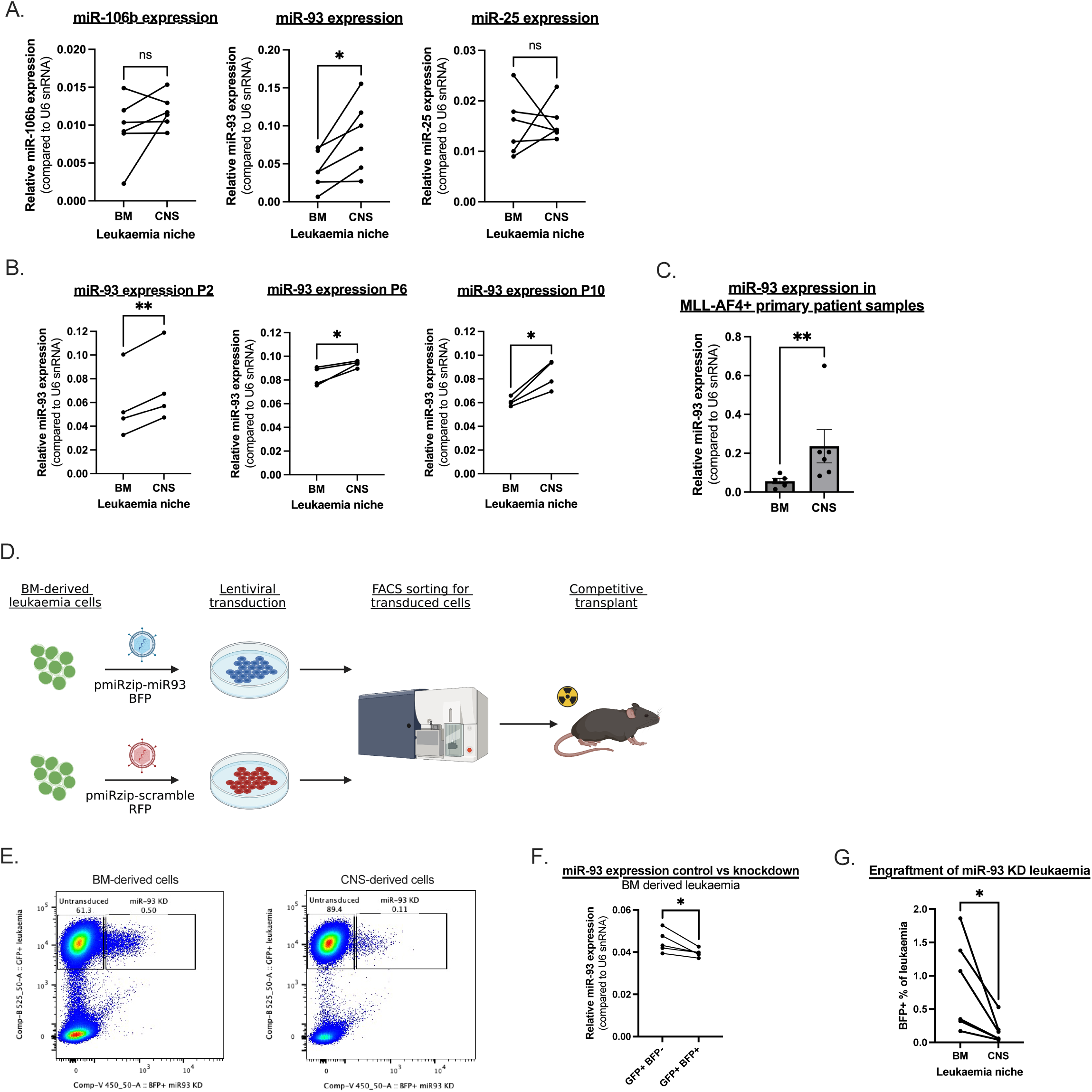
miR-93 expression is upregulated in CNS-derived leukaemia and knockdown impairs engraftment of leukaemia to this niche. (A) Relative expression of miR-106b/25 cluster members in leukaemia cells by niche. MiR expression values are normalised to the expression to U6 snRNA. Increased expression of miR-93 was observed in CNS-derived compared with BM-derived leukaemia cells, with no difference in miR-106b or miR-25 by niche. N=6 paired samples, p values = 0.22, 0.02, 0.86 (miR-106b, miR-93, miR-25 respectively). (B) Validation of niche-specific miR-93 expression in three independent PDX models of KMT2A-AFF1+ infant B-ALL. N=4 paired samples, p = <0.01, 0.04, 0.02 (P2, P6, P10 respectively). (C) Comparison of miR-93 expression between BM and CNS-derived KMT2A-rearrange B-ALL cells from primary patient samples, p <0.01. Paired samples: N =5 BM and N=6 CNS. (D) Experimental overview. (E) Example FACS plots demonstrating the engraftment of GFP+BFP-untransduced leukaemia cells and GFP+BFP+ miR-93 knockdown leukaemia cells in either leukaemia niche. (F) miR-93 expression in FACS-sorted BM-derived GFP+BFP- and GFP+BFP+ cells. Reduced miR-93 expression by GFP+BFP+, i.e. those expressing the miR-93 knockdown vector, N=5 paired samples, p=0.04, fold change=0.88. (G) GFP+BFP+ cell numbers as a percentage of total GFP+ cells in the CNS and BM niche. There was a reduced BFP+ fraction of total leukaemia in the CNS niche compared with the BM niche, N=6 paired samples, p=0.03.

An overall assessment of PI3K pathway activation was performed by comparing the relative phosphorylation status of AKT. There was increased phosphorylated AKT in the CNS-derived infiltrate compared with the BM infiltrate in mice with terminal leukaemia, consistent with increased RTK signalling, decreased pathway antagonism by PTEN, and increased inhibition of the downstream target CDKN1A (Figure 4E). The expected outcome of PI3K/AKT activation would be increased cell survival, growth, and proliferation and represents a novel CNS-specific leukaemia adaptation (Figure 4A).

### miR-93 expression is increased in CNS-derived leukaemia cells across multiple in vivo models of KMT2A-AFF1+ infant B-ALL

The expression of Pten and Cdkn1a is critically regulated by specific miRNA families, which play essential roles in modulating their activity and ensuring proper control over cell cycle progression, growth suppression, and tumour suppression pathways. In silico prediction of upstream miRs using DEGs by niche from miR-128a Kmt2a-AFF1+ LPCs identified overrepresentation of two miR families known to directly regulate Pten and Cdkn1a(N. Li et al. 2017); mmu-miR-22-3p (p=0.004) and mmu-miR-106a-5p/mmu-miR-106b-5p/mmu-miR-17-5p/mmu-miR-20a-5p/mmu-miR-20b-5p/mmu-miR- 6383/mmu-miR-93-5p (p=0.02). The latter family is part of the wider miR-17/92 cluster and its two paralogues, miR-106a/363 and miR-106b/25(Mogilyansky and Rigoutsos 2013). This cluster and its paralogues are also referred to as oncomiR-1. Compared with the other highlighted enriched miRs, there is stronger data to demonstrate direct functional inhibition of Pten and Cdkn1a by the miR-17/92 cluster and therefore, this cluster was selected for further analysis(Sage and Ventura 2011; Ventura et al. 2008). A full list of overrepresented miR families can be found in Supplemental Figure 4.

The expression of miR-106b/25 cluster members was measured in the miR-128a Kmt2a-AFF1+ model by niche based on the strong evidence that supports its role in PI3K regulation and cancer promotion in other tumour types(Shen et al. 2013; Conner et al. 2022; Ohta et al. 2015; W. Yang et al. 2018). Of the three members in this cluster, miR-93 was the only one to be differentially expressed by the niche (Figure 5A, p=0.02, FC=2.1). Its upregulation in the CNS is consistent with decreased expression of Pten and Cdkn1a in CNS-derived cells, which miR- 93 has been shown to inhibit in other studies (Ohta et al. 2015). This miR-93 expression pattern was validated in separate cohorts of KMT2A-AFF1+ infant leukaemia xenograft models from three different patient samples(Figure 5B).

We extended this comparison with B-ALL primary patient samples with KMT2A rearrangements, selected by cytogenetic abnormalities of t(11q23) and/or t(4;11), from a combination of diagnosis and end-of-induction paired BM and CSF samples for those with CNS leukaemia involvement. The age range of these samples was 1-7 years (median age of 2 years), and there was an equal male-to-female ratio, details can be found in Supplemental Table 5. miR-93 was significantly upregulated in CNS-derived samples compared with BM-derived samples (p=0.009, FC=4.1)(Figure 5C).

Upregulation of miR-93 in CNS-derived leukaemia was found in the miR-128a Kmt2a- AFF1+ infant B-ALL model, three KMT2A-AFF1+ infant B-ALL xenograft models and primary patient samples of KMT2A-rearranged B-ALL, validating this niche-specific expression pattern.

### miR-93 knockdown inhibits the development of CNS leukaemia

To understand the *in vivo* functional significance of miR-93 expression, a knockdown and competitive transplant experiment was performed using miR-128a Kmt2a-AFF1+ BM-derived cells transduced with a lentiviral miR-93 knockdown vector (BFP+) or a scramble control (RFP+), which were mixed in a ∼ 8:1 ratio prior to transplant (Figure 5D-E). Mice were culled before the development of terminal leukaemia, on days 18 and 19 post-transplant, to ensure the miR-93 knockdown cells had not been fully eliminated. Cells were harvested from the CNS and BM niche and sorted based on fluorescent protein expression for miR extraction. The sorted cells used for transplant expressed either BFP (miR-93 knockdown) or RFP (miR-scramble control) in addition to GFP (miR-128a Kmt2a-AFF1+), no GFP+BFP-RFP- cells were sorted for transplant. Once mice had developed a leukaemia phenotype following transplant of the sorted transduced cells, analysis of the cell infiltrate showed that no cells expressing RFP were present in either niche, and a GFP+BFP-RFP- population emerged. Either there was contamination with a GFP+BFP-RFP- population, or there was loss/silencing of the miR-scramble control vector in vivo. The latter explanation is favoured as significant contamination would be unlikely since the cell populations were easily distinguishable by flow cytometry and were sorted prior to transplant; an example of a representative

FACS plot can be found in Supplemental Figure 5. GFP+BFP- and GFP+BFP+ cells were sorted for miR extraction to assess the efficacy of the miR-93 knockdown in the GFP+BFP+ cells. Too few GFP+BFP+ cells were present in the CNS niche to sort an adequate number for miR quantification; as a result, this assessment was made only from BM-derived cells. A modest but statistically significant miR-93 knockdown was achieved in GFP+BFP+ cells compared with the GFP+BFP- cells (Figure 5F). Ideally, this comparison would also be made in the CNS-derived cells as the knockdown effect is likely to be larger, given the observation that cells upregulate miR-93 expression in this niche. The modest effect seen in BM-derived cells may be due to low-level miR-93 expression at baseline. Importantly, miR-93 knockdown significantly impairs the engraftment of leukaemia cells in the CNS niche (Figure 5G). It is likely that this impaired engraftment in the CNS niche by miR-93 knockdown is mediated by the resulting reduced PI3K/AKT pathway activity through increased expression of PTEN and CDKN1A.

## Discussion

The aim of this study was to identify novel mechanisms enabling CNS engraftment in infant leukaemia that may explain high rates of CNS involvement in these patients, and that may serve as novel targets for CNS-specific therapies.

We addressed this using a fully murine model of Kmt2a-AFF1+ pro-B infant leukaemia, referred to as the miR-128a Kmt2a-AFF1+ model(Malouf et al. 2021). We began by characterising the LPC populations in the CNS niche, a previously unexplored aspect of the model. We found that both LPC populations (LSK_IL7R and LK/CLP) were present in the CNS niche. When the transplanted LPCs were BM-derived, the BM and CNS leukaemia infiltrates contained the same relative proportions of LSK_IL7R and LK/CLP cells. Transplants of CNS-derived cells, intending to model a CNS to BM relapse event, led to a different overall leukaemia reconstitution phenotype and enriched for a more quiescent cell phenotype (LK/CLPs). We implicate Ube2c as a potential driver of this selection. *Ube2c* encodes a ubiquitin-conjugating enzyme, which is a member of the anaphase-promoting complex and plays a central role in cell cycle progression(Walker et al. 2008). The anticipated effect of increased Ube2c would be to induce cell cycling and, therefore, may be responsible for the increased cell cycling seen in LSK_IL7R LPCs. UBE2C is upregulated in various human non-leukaemia cancers and has recently been studied as an oncogene in hepatocellular carcinoma, where elevated UBE2C expression was associated with proliferation, invasiveness, and therapy resistance(Xiong et al. 2019). Combining the transplant repopulation, the cell cycling and transcriptomic data we speculate that exposure to the CNS niche ‘primes’ LPCs to repopulate a more quiescent leukaemia phenotype on relocation to the BM niche, mirroring a CNS to BM relapse event. Cell cycling differences between LPC populations may be maintained by Ube2c. When relating this to patient relapse dynamics, leukaemia cells that have been exchanged between the CNS and BM niches may become more quiescent and, therefore, more resistant to chemotherapy by virtue of their exposure to the CNS environment. To our knowledge, this is the first published description of the functional difference between BM and CNS-derived LPCs using transplantation assays. If an exchange of leukaemia cells exists between the CNS and peripheral niches in patients, exposure to the CNS niche may induce a slower-cycling leukaemia population and reflect a novel mechanism for acquired chemotherapy resistance.

In this model system, we found transcriptional adaptions to the CNS niche consistent with other published data, specifically adaptations to the low lipid environment with changes in fatty acid and cholesterol metabolism. Upregulation of SCD1 by leukaemia cells in the CNS niche has been identified as a key part of the metabolic rewiring from both microarray analysis of primary patient samples and in xenograft models(Van Der Velden et al. 2016; Savino et al. 2020). An orthologous isoform of SCD1, Scd2, was found to be upregulated in CNS-derived LPCs compared with BM-derived LPCs in our model. Savino et al. identified other DEGs involved in fatty acid metabolism, including the upregulation of the cholesterol transporter ABC1A and downregulation of CPT1A/B, a rate-limiting enzyme in fatty acid oxidation, in CNS B-ALL cells. The miR-128a Kmt2a- AFF1+ B-ALL LPC dataset showed differential regulation of orthologs of these, with upregulation of the Abc1 family member Abca8a and downregulation of Cpt1a in CNS-derived LPCs. These data demonstrate that CNS-derived LPCs from the miR-128a Kmt2a-AFF1+ model are similarly responsive to metabolic stressors present in the CNS niche when compared with other published patient and xenograft datasets. This provides additional validation for this model to be used as a platform for examining CNS leukaemia.

Additional genes, Soat1 and Acot1, involved in fatty acid metabolism were differentially expressed by niche in the miR-128a Kmt2a-AFF1+ model and represent novel CNS leukaemia adaptations. Soat1, which was upregulated in CNS-derived LPCs, is a rate-limiting step in cholesterol metabolism and converts cholesterol to cholesterol-esters, which then form intracellular lipid droplets. SOAT1 upregulation has been explored in multiple cancer subtypes and appears to have additional functions, including activating the PI3K pathway. Acot1, downregulated in CNS-derived LPCs, is also critical to the intracellular regulation of free fatty acids and acyl-CoA esters; however, its role in cancer cell metabolism is not well understood.

Due to the use of an immunocompetent model system, transcriptomic comparisons of LPCs by niche demonstrated differential interactions with immune cells, specifically the differential expression of various immune checkpoints by LPCs by niche. The immune cell landscape in CNS leukaemia is largely unknown and, to our knowledge, these data are the first published comparison of T-cells and macrophages between the CNS and BM niches in B-ALL and highlight some important translatable differences. The genes of multiple immune checkpoints were found to be differentially expressed by niche; this included Ctla4 (the most upregulated gene in CNS-derived LPCs in our dataset), Cd47 and Cd96. Immune checkpoint blockade via anti-CTLA4 monoclonal antibody treatment effectively prevents T cell inactivation through immune tolerance to cancer cell antigens. Less is known about the function of CTLA4 when expressed by malignant cells. However, there are data in other leukaemia subtypes that show CTLA4 expression by leukaemia cells can also suppress T cell function(Do et al. 2019). There is already evidence of clinically significant T cell immune checkpoints expressed by leukaemia cells in KMT2A-AFF1+ infant leukaemia, specifically ICOSLG. A higher ICSOLG expression by leukaemia cells was associated with inferior survival and, using in vitro co-culture assays, ICOSLG induced T-cell differentiation towards immunosuppressive T-regulatory cells(Külp et al. 2022). While Cd96’s regulatory function in the tumour microenvironment is less well-understood than that of Ctla4, it can mediate an inhibitory signal for T and NK cell effector function and proliferation through interactions with CD155 expressed on tumour cells. Upregulation of *CD96* has been linked to the failure of immune checkpoint blockade therapy(Feng et al. 2023).

CD47 binds to SIRPα on the surface of macrophages to deliver an antiphagocytic signal. The upregulation of CD47 by cancer cells has been identified as a critical immune escape mechanism across haematological and non-haematological malignancies, although not ALL, and several CD47-directed monoclonal antibodies are currently being assessed in clinical trials(Z. Jiang et al. 2021).

It is, therefore, likely that niche-dependent expression of multiple immune checkpoints by leukaemia cells as we observed here will impact immune-mediated leukaemia clearance in the CNS niche. This may be due to environmental factors, such as stromal secretion of immunosuppressive cytokines, leukaemia-induced immunosuppression or chronic antigen exposure due to peripheral T cell priming and CNS trafficking. Indeed, our data suggest impaired T cell functioning and a less phagocytic macrophage phenotype in the CNS leukaemia niche compared with the BM. We speculate that this may provide a more permissive immune environment for leukaemia. There is inconsistent data regarding T cell redirection therapies, including blinatumomab and CAR-T cells, for CNS B-ALL; however, some reports suggest reduced efficacy compared with BM-only disease(Brown 2021; Jacoby et al. 2022). Linking these clinical outcomes to our data, we also hypothesise that immunotherapy failure in CNS leukaemia may be due to environmentally driven factors, such as metabolic adaptations, limiting immunotherapy-induced immune responses.

One of the ways cancer-associated macrophages have been shown to support cancer cells is through the secretion of soluble factors in the niche, activating growth factor pathways, including PI3K(D. Li et al. 2020; Mafi et al. 2021). As a result, we focused on a novel differential gene expression pattern between CNS and BM-derived LPCs that regulates PI3K signalling. This line of investigation is further supported as there are existing data that PI3K inhibition reduces CNS B-ALL progression in murine xenograft models, although not previously explored in KMT2A-rearranged or infant ALL(Ridge et al. 2021; Yao et al. 2018). Transcriptomic and protein level validation data suggested the activation of PI3K signalling through the upregulation of c-MET, and the downregulation of PTEN and CDKN1A may act as a pro-survival mechanism for leukaemia in the CNS niche. This suggests niche-specific differential regulation of growth factor pathways, specifically the PI3K pathway, in LPCs. Dysregulation of the PI3K pathway involving aberrant PTEN expression is common across a multitude of cancer subtypes; its activation is associated with tumour invasiveness and metastatic ability(J. Yang et al. 2019; Kim and Guan 2019). This context highlights the modulation of the PI3K pathway as a strong candidate for CNS-specific survival adaptations in KMT2A-AFF1+ infant leukaemia and provides additional data to strengthen its case as a CNS-leukaemia therapeutic target.

To build on these mechanistic insights we explored higher level regulatory mechanisms as potential biomarkers of CNS leukaemia. We linked the RNA expression data to a known regulator of Pten and Cdkn1a expression – miR-93. The miR-106b/25 cluster, which includes miR-93, has been explored as an emerging oncomir across many cancer subtypes with multiple molecular mechanisms, including activation of the PI3K pathway(Mehlich, Garbicz, and Włodarski 2018). However, this miR cluster has received little attention in the pathogenesis of haematological malignancies. We demonstrated the reproducibility of miR-93 upregulation in CNS compared with BM-derived KMT2Ar B-ALL samples. This included the miR-128a Kmt2a-AFF1+ model, three independent xenograft mouse models of KMT2A-AFF1+ infant leukaemia and KMT2A- rearranged primary patient B-ALL samples. Furthermore, a knockdown of miR-93 in the miR-128a Kmt2a-AFF1+ model demonstrated impaired CNS leukaemia engraftment.

The relevance of these findings is twofold. Firstly, we have described novel gene expression and a possible mRNA-miR regulatory network specific to CNS leukaemia in KMT2A-rearranged B-ALL. This may provide a biological mechanism for the high level of CNS involvement in this patient population, and it opens an avenue to novel targeted therapeutics. Secondly, we have reliably demonstrated the upregulation of miR-93 in CNS KMT2A-AFF1+ B-ALL, which could directly impact clinical biomarker development. CNS leukaemia biomarkers are lacking, and cerebrospinal fluid-free miRs are one of the more promising candidates under investigation(Thastrup et al. 2022; Egyed et al. 2020).

## Method

### Mice and transplantation experiments

All animal work was approved by the University of Edinburgh’s Bioresearch and Veterinary Services and undertaken in accordance with institutional and UK Home Office guidance. Details of the mice used for RNA sequencing and LPC comparisons can be found in Supplemental Table 4.

miR-128a Kmt2a-AFF1+ LPCs (GFP+) from mice that developed terminal leukaemia were serially transplanted into sublethally irradiated (2 doses of 4.6 Gy, 3 hours apart, with a split adaptor) 8–12 week-old recipients (CD45.1/2 C57BL/6NCrl) through tail-vein injection. 2,000-150,000 cells per transplant suspended in 2% fetal calf serum (FCS) in phosphate-buffered saline (PBS) were used. Mice were administered antibiotics in their drinking water (0.1mg/mL enrofloxacin; 10% Baytril (Bayer)) post-transplant for a period of 28 days.

For PDX models, three samples were used for xenografts: P2, P6, and P10, which were BM aspirates from KMT2a-AFF1+ infant B-ALL patients at diagnosis, clinical characteristics are described in Supplemental Table 5. 100,000 cells were transplanted by tail vein into non-irradiated NSG mice aged 8-12 weeks old. Mice were administered antibiotics in their drinking water (0.1mg/mL enrofloxacin; 10% Baytril (Bayer)) for 3 days after transplantation.

At the end of the experiment, when the mice developed symptoms of leukaemia according to a pre-defined scoring system (including diminished activity levels and poor body condition), they were culled by CO2 asphyxiation.

### Isolation of leukaemia cells

For murine BM LPCs, femurs and tibias of mice that developed leukaemia were dissected and crushed in ice-cold PBS. For CNS LPCs, heads were recovered and surrounding soft tissue was removed. The base of the skill was removed, brain was then removed intact and placed in a 15mL Falcon tube containing 10mL of ice-cold PBS.

Using a 1mL syringe plunger and Pasteur pipette, the meninges/skull vault was scraped, flushed with PBS and collected in the Falcon tube containing the brain. The Falcon tube was then gently vortexed for 2 minutes to dislodge cells from the outside of the brain. The collected cells were pelleted and resuspended in Flow Cytometry Staining Buffer (ThermoFisher, 00-4222-26). Cells were stained with the following antibodies for the detection of LPCs (LSK_IL7R and LK/CLP): APC-eF780 anti-mouse CD45.2 antibody (clone 104, eBioscience, 47-0454-82), APC anti-mouse CD3ε antibody (clone I45-2C11, Biolegend, 100312), APC anti-mouse TER-119 antibody (clone TER119, Biolegend, 116212), APC anti-mouse F4/80 antibody (clone BM8, Biolegend, 123116), APC anti-mouse Nk1.1 antibody (clone PK136, Biolegend, 108709), APC anti-mouse Ly- 6G/Ly-6C (Gr-1) antibody (clone RB6-8C5, Biolegend, 108412), APC anti-mouse/human B220 antibody (clone RA3-6B2, Biolegend, 103211), APC anti-mouse CD19 antibody (clone 6D5, Biolegend,115511), Brilliant Violet 421 anti-mouse c-Kit (CD117) antibody (clone 2B8, Biolegend, 105827), PE/Cy7 anti-mouse Ly-6A/E (Sca-1) antibody (clone E13-161.7, Biolegend, 122513), PE anti-mouse IL7-R (CD127) antibody, eBioscience (clone A7T34, ThermoFisher, 12-1271-82), biotin anti-mouse Flt3 (CD135 antibody), eBioscience (clone A2F10, ThermoFisher, 13-1351-81). The stained cells were incubated on ice, washed twice and resuspended in Flow Cytometry Staining Buffer with the viability dye SYTOXAADvanced (ThermoFisher, S10274). Cells were sorted using an BD FACSAria II (BD Biosciences). LPCs were sorted using the following strategy: live cells, CD45.2 positive, GFP positive, lineage negative (CD3ɛ, Ter119, F4/80, Nk1.1, Gr1, B220 and CD19), c-Kit positive, Sca-1 negative and Sca-1 positive IL7-R positive. Xenograft-derived human leukaemia cells were isolated based on viability (using DAPI) and by human CD45 (APC, clone 2D1, Biolegend, 368511) positivity and murine CD45 (PE-Cy7, clone 30-F11, Biolegend, 103113) negativity. Flow cytometry standard (*.fcs) files generated from each experiment were imported into FlowJo v10.8 Software (BD Life Sciences) for analysis.

### Histology and immunofluorescence staining

Imaging of the meningeal leukaemia infiltrate was performed on paraffin-embedded, formalin-fixed and decalcified heads of mice that develop leukaemia. Intact heads stripped of soft tissue were fixed in CellStor Pots (10% formalin, BAF-600-08A, CellPath) for a minimum of 7 days and then decalcified in 200mL of EDTA decalcification solution (188mM EDTA solution with 4% formaldehyde in water) for three weeks. Slides were imaged using a Zeiss Axioscan Slide Scanner.

Immunofluorescence staining was performed on 50,000-100,000 cells concentrated on a slide using a Cytospin 4 Cytocentrifuge (Shandon Thermo). Cell spots were fixed in methanol for 5 minutes and then washed in PBS twice. A PAP pen (Liquid Blocker Super PAP Pen, Diado Sangyo) created a hydrophobic barrier around each cell spot on the slides. A volume of 200μL blocking buffer (PBS + 0.05% Tween + 1% BSA, supplemented with 5% donkey serum from Jackson Immunoresearch, 017-000-121-JIR) was added to the cell spot and left to incubate for 1 hour at room temperature (RT). Next, the unconjugated primary anti-c-kit antibody (raised in rabbit, clone EPR24708-25, ab273119, Abcam) was diluted 1:200 in blocking buffer and applied to the cell spot for 1 hour at RT in the dark. Following this, the slides underwent three washes with PBS containing 0.05% Tween, each for 5 minutes. The primary anti-F4/80 antibody, conjugated with Alexa Fluor 647 (raised in rat, clone F4/80, ab204467, Abcam) and diluted 1:200 in blocking buffer, along with the secondary anti-rabbit IgG (H+L) conjugated with Alexa Fluor 488 (raised in donkey, polyclonal, A-21206, Thermo Fisher) diluted 1:500 in blocking buffer, were incubated on the cell spot for 1 hour at RT in the dark. The slides were then washed three times in PBS with 0.05% Tween. DAPI nuclear staining (diluted 1:2500, Life Tech, D1306) was performed by incubating the cell spot for 5 minutes at RT in the dark, followed by three washes in PBS with 0.05% Tween. The slides were subsequently dehydrated using ethanol. Finally, slides were mounted with ProLong Glass Antifade Mountant (P36980, Invitrogen), and coverslips were placed on top, after which they were imaged using an Operetta high-content microscope.

### RNA sequencing and analysis

An overview of the workflow is presented in Supplemental Figure 3. RNA was extracted from freshly sorted LPCs using the Quick-RNA Microprep Kit (Zymo, R1050). cDNA library preparation was performed using SMARTer Stranded Total RNA-Seq Kit v3 – Pico Input Mammalian (TaKaRa, 634487) followed by 100bp paired-end sequencing with a NextSeq 2000 instrument (Illumina) by Glasgow Polyomics. Raw reads underwent quality control analysis using the FastQC tool(Andrews 2010). The Cogent NGS Analysis Pipeline v2.0 (TaKaRa) in the command line was used for data preprocessing. This function performs read trimming using Cutadapt and sequence alignment to a reference genome using STAR (Spliced Transcripts Alignment to a Reference)(Martin 2011; Dobin et al. 2013). Genome Reference Consortium Mouse Build 39 (GRCm39) was used as the reference genome. SummarizedExperiment objects containing count matrices were created using the GenomicAlignments package and the aligned BAM files. SummarizedExperiment objects were transformed into DESeqDataSet objects, prefiltered, normalised and a results table created using the DESeq2 package (Lawrence et al. 2013; Love, Huber, and Anders 2014). Additional steps were taken when comparing human and murine data sets: biomaRt was used to convert human genes to murine orthologs and batch effects accounted for using Limma(Durinck et al. 2009; Ritchie et al. 2015). DESeqDataSet objects were transformed using variance stabilising transformation and underwent dimensionality reduction by PCA using DESeq2. Gene ontologies were identified using clusterProfiler(Wu et al. 2021).

The RNA-sequencing data has been deposited at GEO (GSE280505). The other RNA-sequencing datasets used in this project were from Symeonidou et al. (GEO: GSE167234) and Andersson et al. (EGA: EGAS00001000246).

### Flow cytometry and cell cycle analysis

Flow cytometry for immune cell populations used predominantly extracellular staining only, except for panels assessing T regulatory cells and cell cycle which required intracellular and extracellular antibody staining. For extracellular-only staining, cells were pelleted, resuspended in an appropriate antibody cocktail, and incubated in the dark on ice for 20 minutes. Cells were washed twice using ice-cold PBS and resuspended in FACS buffer (2% FBC in PBS with 1% penicillin/streptomycin (Sigma-Aldrich)) containing viability dye.

For intracellular and extracellular staining to detect T regulatory cells, the True-Nuclear Transcription Factor Buffer Set (Biolegend, 424401) was used according to manufacturer instructions. For *in vivo* cell cycle analysis, Phase-Flow Alexa Fluor 647 BrdU Kit (Biolegend, 370706) was used according to the manufacturer’s instructions. Bromodeoxyuridine (1mg) was administered to mice with established leukaemia by intraperitoneal injection 90 mins before retrieving the cells.

The following antibodies and viability stains were used. For T cell subsets: APC-eF780 anti-mouse CD45.1 antibody (clone A20, eBioscience, 47-0453-82), FITC anti-mouse CD45.2 antibody (clone 104, Biolegend, 109807), BV711 anti-mouse CD3 antibody (clone I45-2C11, Biolegend, 100349), BV421 anti-mouse CD4 antibody (clone GK1.5, Biolegend, 100437), PerCP-Cyanine5.5 anti-mouse CD8 antibody (clone 53-6.7, eBioscience, 45-0081-82), PE anti-mouse IL2-Ra (CD25) antibody (clone A7R34, eBioscience, 12-1271-82), AF647 anti-mouse Foxp3 antibody (clone MF-14, Biolegend, 126408), Fixable Viability Dye eFluor 455UV (eBioscience, 65-0868-14). For T cell activity markers: APC-eF780 anti-mouse CD45.1 antibody (clone A20, eBioscience, 47- 0453-82), BV711 anti-mouse CD3 antibody (clone I45-2C11, Biolegend, 100349), APC anti-mouse CD4 antibody (clone GK1.5, Biolegend, 100411), PerCP-Cyanine5.5 anti-mouse CD8 antibody (clone 53-6.7, eBioscience, 45-0081-82), PE-Cy7 anti-mouse CTLA4 (CD152) antibody (clone UC10-4B9, Biolegend, 106313), PE anti-mouse PD-1 (CD279) antibody (clone RMP1-30, Biolegend, 109103), PE anti-mouse CD69 antibody (clone H1.2F3, Biolegend, 104507), 4,6-diamidino-2-phenylindole (DAPI) (Life Tech, D1306). For NK cells: APC-eF780 anti-mouse CD45.1 antibody (clone A20, eBioscience, 47-0453-82), PeCy7 anti-mouse Nk1.1 antibody (clone PK136, Biolegend, 108713), BV607 anti-mouse NKp46 (CD335) antibody (clone 29A1.4, Biolegend, 137619), PE anti-mouse CD69 antibody (clone H1.2F3, Biolegend, 104507), DAPI (Life Tech, D1306). For macrophages: PE anti-mouse CD45.1 antibody (cloneA20, eBiosciences, 12-0543-83), BV711 anti-mouse Ly6C-G antibody (clone RB6-8CA, Biolegend, 108443), APC anti-mouse F4/80 antibody (clone BM8, Biolegend, 123115), PeCy7 anti-mouse I-A/I-E antibody (clone M5/114.15.2, Biolegend, 107629), BUV395 anti-mouse CD11b antibody (clone M1/70, BD Biosciences, 565976), DAPI (Life Tech, D1306). For cell cycle analysis: PE anti-mouse CD45.1 antibody (clone A20, eBioscience, 12-0543-83), FITC anti-mouse CD45.2 antibody (clone 104, Biolegend, 109807), BV785 anti-mouse c-Kit (CD117) antibody (clone 2B8, Biolegend, 105841), Pacific Blue anti-mouse Ly-6A/E (Sca-1) antibody (clone E13-161.7, Biolegend, 122519), AF647 anti-BrdU antibody (Biolegend, 370706), 7AAD (Biolegend, 370706).

Flow cytometry was performed using a 5-laser BD LSRFortessa and a 3-laser Cytek Aurora. Flow cytometry standard (*.fcs) files generated from each experiment were imported into FlowJo v10.8 Software (BD Life Sciences) for analysis.

### Western blot

Cell lysis and total protein extraction were performed using radioimmunoprecipitation assay (RIPA) buffer. The RIPA buffer contains 50 mM sodium chloride (CHE3326, Scientific Labs), 1% Triton X-100 (T8787, Sigma-Aldrich), 0.5% sodium deoxycholate (30970, Sigma-Aldrich), 0.1% sodium dodecyl sulfate (SDS) (S/P530/53, Fisher Scientific), 50mM trisaminomethane (Tris) (Trizma base, T6066, Sigma-Aldrich) pH 8.0 in water. This buffer was supplemented with a protease inhibitor (cOmplete Protease Inhibitor Cocktail, Roche, 4693116001) (10mL RIPA buffer per one tablet of protease tablet). ∼5x10^6^ LPCs were used for protein extraction. DNA was fragmented using needle shearing by passing each sample through a 19G needle (BD, 301500) and 1mL syringe (BD, 300013) up to 10 times. A colourimetric assay (DC Protein Assay Kit II, BioRad, 5000112) was used to quantify proteins using a BMGLabtech FLUOstar Omega microplate reader.

1mm hand-cast gels were made using the Mini-PROTEAN Tetra Cell Handcasting kit (BioRad, 1653370). Two concentrations of acrylamide PAGE gels were used. Separation gel: 12% acrylamide (37.5:1) (BioRad, 1610158), 625mM Tris pH 8.8, 0.17% SDS, 0.08% tetramethylethylenediamine (TEMED) (Sigma-Aldrich, T9281), 0.4% ammonium persulfate (APS) (Fisher Scientific, A/P460?46) in water. Stacking gel: 4% acrylamide (37.5:1), 120 mMTris pH 6.8, 0.1% SDS, 0.01% TEMED, 0.4% APS in water.

10μg of extracted protein was added to 20μL of 2X Laemmli Sample Buffer (BioRad, 161-0737) and the final volume was adjusted to 40μL with water. Protein samples were then heat denatured. Protein migration proceeded with a Tris-Glycine-SDS running buffer (25mMTris, 190mMglycine (G/0800/60, Fisher Scientific), 0.1% SDS in water) alongside a Precision Plus Protein Dual Color Standard (BioRad, 161-0374). Migration in the stacking gel took place at 80V and in the separation gel at 150V.

Proteins were transferred from the separation PAGE gel to a polyvinylidene difluoride (PVDF) membrane (BioRad, 1620177) using the wet transfer method. Transfer occurred using a transfer buffer (25 mM Tris base, 190 mM glycine, 10% methanol in water) at 100V over 1 hour in ice.

Membranes containing the transferred proteins were blocked with Blotto A (5% skimmed milk (84615, VWR Chemicals) in Tris-buffered saline with 0.05% Tween 20 (TBST: 50mM Tris, 150mMNaCl in water) for 1 hour at RT with shaking. Membranes were incubated with primary antibodies diluted in Blotto A overnight at 4°C with shaking: anti-mouse beta-actin antibody (clone AC-74, Sigma-Aldrich, A2228), anti-rabbit AKT antibody (polyclonal, Cell Signalling Technology, 9272), anti-rabbit phosphor-AKT (Ser473) antibody (polyclonal, Cell Signalling Technology, 9271), anti-rabbit CDKN1a antibody (clone EPR18021, Abcam, ab188224), anti-rabbit PTEN antibody (clone 138G6, Cell Signalling Technology, 9559). Following washing, membranes were incubated with 10mL of Blotto A containing 1:5000 horseradish peroxidase (HRP)- conjugated secondary antibody at RT for 1 hour with shaking: goat anti-mouse IgG (H+L) secondary antibody (Invitrogen, A16066), goat anti-rabbit IgG (H+L) secondary antibody (Invitrogen, A16096). Membranes were then washed three times for 5 minutes with TBST and one time for 5 minutes with TBS.

Membranes were incubated with 2mL 1:1 Clarity Western Peroxide Reagent and Clarity Western Luminol/Enhancer Reagent (Clarity Western Enhanced Chemiluminescence (ECL) Substrate, BioRad, 1705061) for 1 minute. The resulting chemiluminescence was imaged using a ChemiDoc with Blot/UV/Stain-Free Sample Tray (BioRad, 17001401).

Images were imported into Image Lab Software (version 6.1, BioRad, SOFT-LIT-170- 9690-ILSPC-V-6-1) for analysis.

### MicroRNA enrichment analysis

Mienturnet (MicroRNA ENrichment TURned NETwork), using the TargetScan and miRTarBase databases, was used for *in silico* prediction of gene-miR regulatory networks(Licursi et al. 2019).

### MicroRNA quantification by PCR

MiR were extracted from leukaemia cells using the miRNeasy Mini Kit (Qiagen, 217044). MiRs of interest were quantified by real-time quantitative PCR (RT-qPCR) using the TaqMan miR Reverse Transcription Kit and TaqMan Universal Master Mix II, no UNG (Thermo Fisher, 4366596 and 4440040) using a QuantStudio 7 Flex instrument (Thermo Fisher) according to the manufacturer’s instructions. The following TaqMan MicroRNA Assays (Thermo Fisher, 4427975) were used: U6 (assay id: 001973), miR-25 (assay id: 00403), miR-93 (assay id: 001090), miR-106b (assay id: 000442), miR-128a (assay id: 002216).

Primary patient samples and data used in this study were provided by VIVO Biobank, supported by Cancer Research UK & Blood Cancer UK (Grant no. CRCPSC- Dec21\100003). One BM sample was excluded from the final analysis due to extremely poor post-thaw viability of 2%. The sample details are described in Supplemental Figure 5.

### Lentiviral production and transduction

Two lentiviral vectors were used: miRZip anti-miR-93 Expression Lentivector customised to express BFP (MZIP93-PA-1, System Biosciences) and miRZip Scramble Lentivector customised to express RFP (MZIP000-PA-1, System Biosciences). pMD2.G (12259, Addgene) and psPAX2 (12260, Addgene) were used for enveloping and packing these vectors. Heat shock transformation and ampicillin selection was used to clone and propagate DNA vectors using One Shot Stbl3 Chemically Competent E. Coli (C737303, Thermo Fisher). This was followed by liquid culture, and plasmid purification using Plasmid Midiprep (12643, Qiagen).

Either miRZip lentivectors, pMD2.G and psPAX2 were mixed with 1mg/ml high molecular weight polyethyleneimine (408727, Sigma-Aldrich) in serum-free DMEM (41966029, Gibco). The mix was incubated at RT for 15 mins and added to HEK293T cells and cultured in DMEM,10% fetal bovine serum (FBS) (heat-inactivated, SV30180.03, Fisher Scientific), 1% penicillin/streptomycin (P/S) (Sigma-Aldrich), and 1% L-glutamine (25030081, Gibco), following culture overnight the media was replaced. Supernatant containing the lentivirus was collected after a further 48 hours of culture and filtered through a Millex-hV 0.45μmPVDF 33mmGamma Sterilized filter (SLHVM33RS, Millipore) prior to use.

Virus-containing supernatant was added to untreated plates coated in RetroNectin Recombinant Human Fibronectin Fragments (T100A, TaKarRa). Leukaemia cells were then added and cultured overnight in StemPro-34 Serum-FreeMedia (10639011, Thermo Fisher) supplemented with 100ng/mL SCF (315-14, Peprotech), 100ng/mL TPO (250-03, Peprotech), 50ng/mL Flt3-Ligand (250-31L, Peprotech), and 100ng/mL IL7 (217-17, Peprotech). Harvested cells were FAC-sorted for viability and either GFP+BFP+ or GFP+RFP+ prior to transplantation. There was a difference in the yield of transduced cells that resulted in each transplant recipient receiving 68,000 GFP+BFP+ and 8,200 GFP+RFP+ cells by tail vein injection.

### Statistical analysis

The raw data were imported into GraphPad Prism 9 for statistical analysis and graph generation. Normality was assessed for each dataset before applying either a paired or unpaired T-test or a non-parametric alternative (Wilcoxon matched-pairs signed rank test or Mann-Whitney test, respectively). For comparisons involving three or more groups, two-way ANOVAs were conducted. Error bars in the graphs represent the standard error of the mean. P-values are indicated in graphs with asterisks marking levels: p < 0.05 (*), p < 0.01( **), and p < 0.001 (***).

## Authorship contribution

A.D. designed the study, performed experiments, analysed results and wrote the manuscript. C.M. and L.N. assisted with performing experiments. N.A.B and O.P.S. provided patient material. C.H. helped design the study and provided supervision to the study. K.O. conceived and supervised the study and wrote the manuscript. All authors approved the submission of the final version of the manuscript.

## Acknowledgements

We are tremendously grateful to the patients and their parents for donating the samples used in this study including those provided by VIVO Biobank. We would like to thank the core services at the Centre for Regenerative Medicine, in particular the Flow Cytometry Service, Animal Facility and Imaging Facility. From these facilities we would like to thank and acknowledge the support of Fiona Rossi, Claire Cryer, Andrea Corsinotti, James Todd, Allan Booth, Jacek Mendrychowski, Jaimie Kelly, Monika Struzik, Matthieu Vermeren and Justyna Cholewa-Waclaw. We would like to thank the Institute for Regeneration and Repair’s Histology Facility including Melanie McMillan. We would like to thank the Glasgow Polyomics facility including Julie Galbraith and Graham Hamilton. This work was funded by a Cancer Research UK Clinical Research Fellowship (C60521/A31315 to AD) and Cancer Research UK Programme Foundation Award (C57303/A23581 to KO and DRCPFA-Nov21\100001 to CH).

## Conflict of Interests

The authors have no conflicts of interest to disclose.

## Grant support

Cancer Research UK Clinical Research Fellowship (C60521/A31315 to AD) and Cancer Research UK Programme Foundation Award (C57303/A23581 to KO, and DRCPFA-Nov21\100001 to CH).

---

**Supplemental Figure 1. Leukaemia phenotype comparison by type of BM-derived LPC transplanted.** (A) No difference in survival by BM-derived LPC type transplanted in the miR-128a Kmt2a-AFF1+ model. Number of replicates: LSK_IL7R transplanted N=13, LK/CLP transplanted N=15. (B) No difference in spleen and liver weights of mice that developed terminal leukaemia by BM-derived LPC type transplanted. Number of replicates: LSK_IL7R transplanted N=9, LK/CLP transplanted N=12. (C) PB counts and representative blood film image of mice that developed terminal leukaemia with no differences by BM-derived LPC type transplanted. Number of replicates: LSK_IL7R transplanted N=9, LK/CLP transplanted N=12. (D) Equal expression of myeloid and lymphoid cell surface antigens by BM-derived LPC type transplanted. Number of replicates: LSK_IL7R transplanted N=4, LK/CLP transplanted N=4. (E) No difference in transplanted cell engraftment by BM-derived LPC type transplanted; mean of 96% engraftment for LSK_IL7R cells transplanted and 93% for LK/CLP cells transplanted. Number of replicates: LSK_IL7R transplanted N=7, LK/CLP transplanted N=5. Image of a representative cytospin of bone marrow infiltrate showing a monomorphic blasts population. (F) Both LPCs reconstituted equal proportions of LSK_IL7R and LK/CLP in the BM of mice that developed terminal leukaemia. Number of replicates: LSK_IL7R transplanted N=7, LK/CLP transplanted N=5.

**Supplemental Figure 2. Leukaemia phenotype comparison by type of CNS-derived LPC transplanted and by source of LPC.** (A) No difference in survival by CNS-derived LPC type transplanted in the miR-128a Kmt2a-AFF1+ model. Number of replicates: CNS-derived LSK_IL7R transplanted N=8, CNS-derived LK/CLP transplanted N=8. (B) No difference in survival by source of LPC type transplanted. Number of replicates: CNS-derived LPCs transplants: LSK_IL7R N=8, LK/CLP N=8 and BM-derived LPCs transplants: LSK_IL7R N=13, LK/CLP N=15. (C) No difference in spleen and liver weights of mice who developed terminal leukaemia by CNS-derived LPC type transplanted. Number of replicates: CNS-derived LSK_IL7R transplanted N=8, CNS-derived LK/CLP transplanted N=7. (D) No difference in spleen and liver weights of mice that developed terminal leukaemia by source of LPC type transplanted. Number of replicates: CNS-derived LPCs transplants: LSK_IL7R N=8, LK/CLP N=7 and BM-derived LPCs transplants: LSK_IL7R N=9, LK/CLP N=12. (E) No difference in PB counts by type of CNS-derived LPC or by the source of LPC. Number of replicates: CNS-derived LPCs: LSK_IL7R N=5, LK/CLP N=6 and BM-derived LPCs: LSK_IL7R N=6, LK/CLP N=7. Number of replicates: CNS-derived LPCs: LSK_IL7R N=4, LK/CLP N=4 and BM-derived LPCs: LSK_IL7R N=3, LK/CLP N=3. (B, D-E) For these comparisons, data from LSK_IL7R and LK/CLP cell transplants were pooled, comparing quaternary transplants from BM-derived LPCs with CNS-derived LPCs (quinary recipients).

**Supplemental Figure 3. The miR-128a overexpression Kmt2a-AFF1+ model of infant B-ALL is representative of human disease by transcriptomic comparisons.** (A) PCA showing preferential clustering of miR-128a Kmt2a-AFF1+ model LPCs (“miR128a_model”) with infant KMT2A-AFF1+ B-ALL patient samples (“infant_ALL”) compared with paediatric KMT2A-AFF1+ B-ALL patient samples (“paediatric_ALL”), by differentially expressed genes between infant and patient KMT2A-AFF1+ B-ALL samples. PCA (B) and heatmap (C) showing clustering of miR-128a Kmt2a-AFF1+ LPCs (BM_proBALL: LK and LSK_IL7R) with wild-type FL LMPPs (FL_WT LMPP). The heat map shows DEGs between FL and adult BM LMPPs (BM_WT LMPP), p.adj<0.05, ordered by log2FC.

**Supplemental Figure 4. RNA sequencing workflow.** The figure summarises the RNA sequencing workflow. The table summarises the range of FACS-sorted cell numbers used for RNA extraction.

**Supplemental Figure 5. List of overrepresented miR families.** List of significantly overrepresented miR families, by p<0.05, based on inferred regulation of DEGs between CNS and BM-derived LPCs from the miR-128a Kmt2a- AFF1+ model. Overrepresented miR families with Pten and Cdkn1a as targets are highlighted. The asterisk (*) indicates the transplant stage of mice for the comparisons of LPC repopulation dynamics by source, i.e. comparing initial transplants from BM-derived LPCs with CNS-derived LPCs (subsequent transplant).

**Supplemental Figure 6. Representative FACS plot comparing BM and CNS engraftment of miR-93 knockdown of miR-128a Kmt2a-AFF1+ LPCs.** (A-B) FACS sorting data of cells used for competitive miR-93 knockdown versus control transplant. The data demonstrate miR-128a Kmt2a-AFF1+ cell transduction with miRZip anti-miR-93 BFP Expression Lentivector (A) and control miRZip Scramble RFP Expression Lentivector (B). (C) Example FACS data from BM and CNS samples of mice following transplantation of miR-128a Kmt2a-AFF1+ miR-93 knockdown and control miRZip scramble control cells. miR-128a Kmt2a-AFF1+ miR-93 knockdown (GFP+ BFP+) are shown to be engrafted in both niches, but miR-128a Kmt2a-AFF1+ miRZip scramble control cells (GFP+ RFP+) are not present in either.

**Supplemental Table 1. Transplant details of mice used for RNA sequencing and LPC comparisons.**

**Supplemental Table 2. Differentially expressed genes between BM-derived LSK_IL7R and LK/CLP cells.**

**Supplemental Table 3. Differentially expressed genes between CNS-derived LSK_IL7R and LK/CLP cells.**

**Supplemental Table 4. Differentially expressed genes between CNS-derived and BM-derived LPCs.**

**Supplemental Table 5. Characteristics of patient samples.**

## Notes

### Competing Interest Statement

The authors have declared no competing interest.

