## Supplemental figures and tables for "Leukaemia cell intrinsic and extrinsic factors cooperate to facilitate the survival and proliferation of KMT2A-rearranged B-ALL in the CNS niche"

Supplementary Figure 1

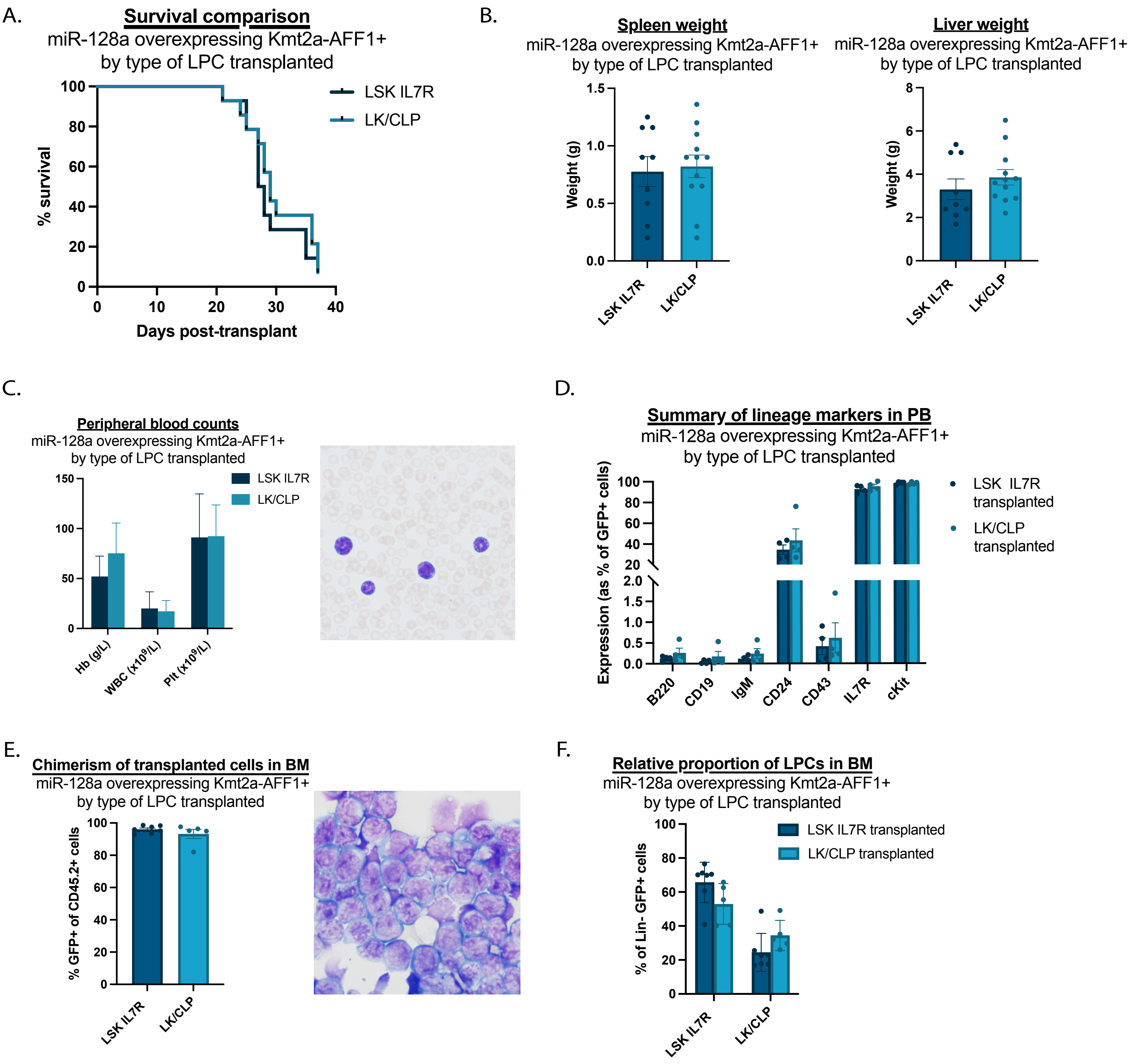

Supplementary Figure 2

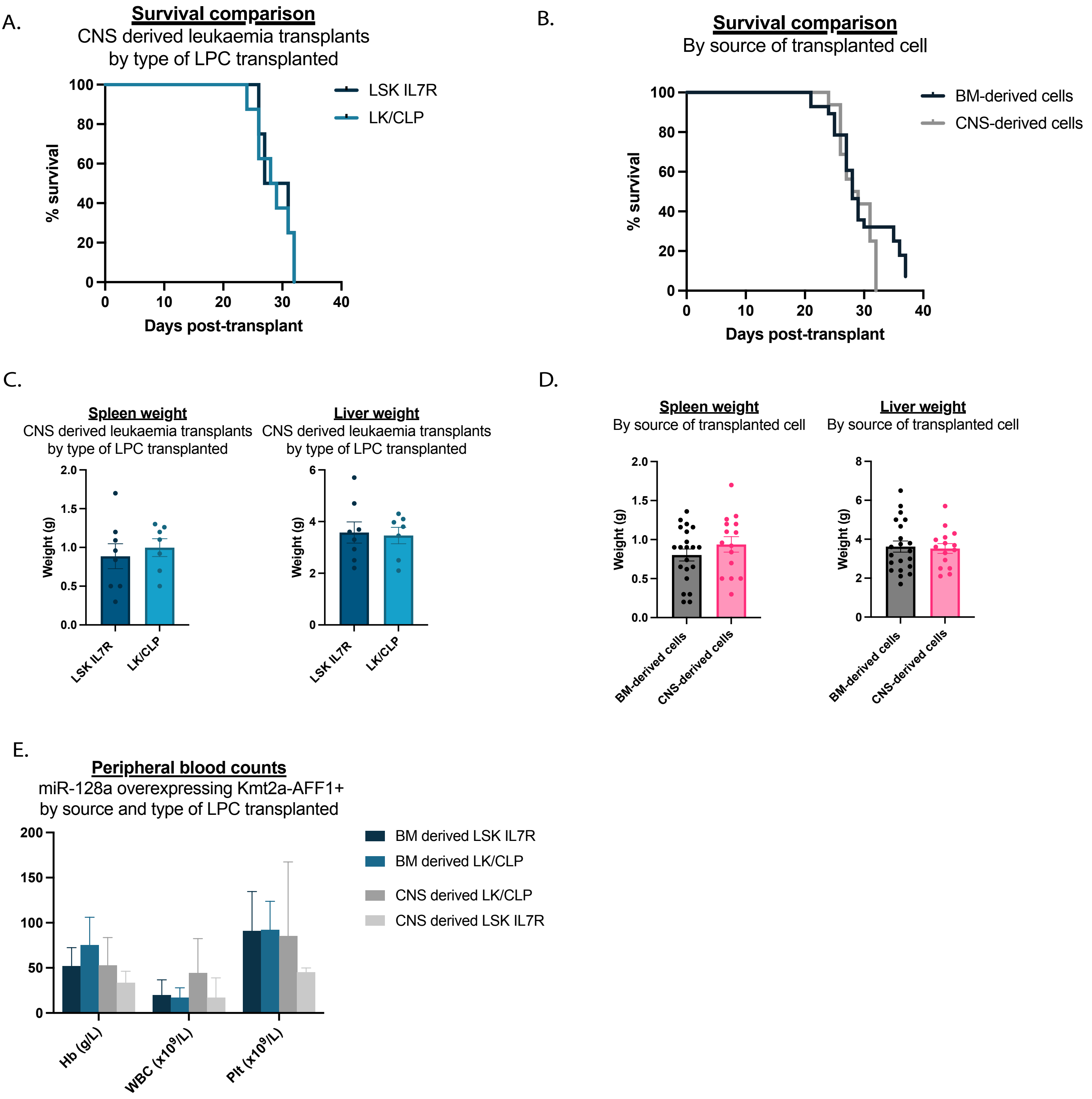

Supplementary Figure 3

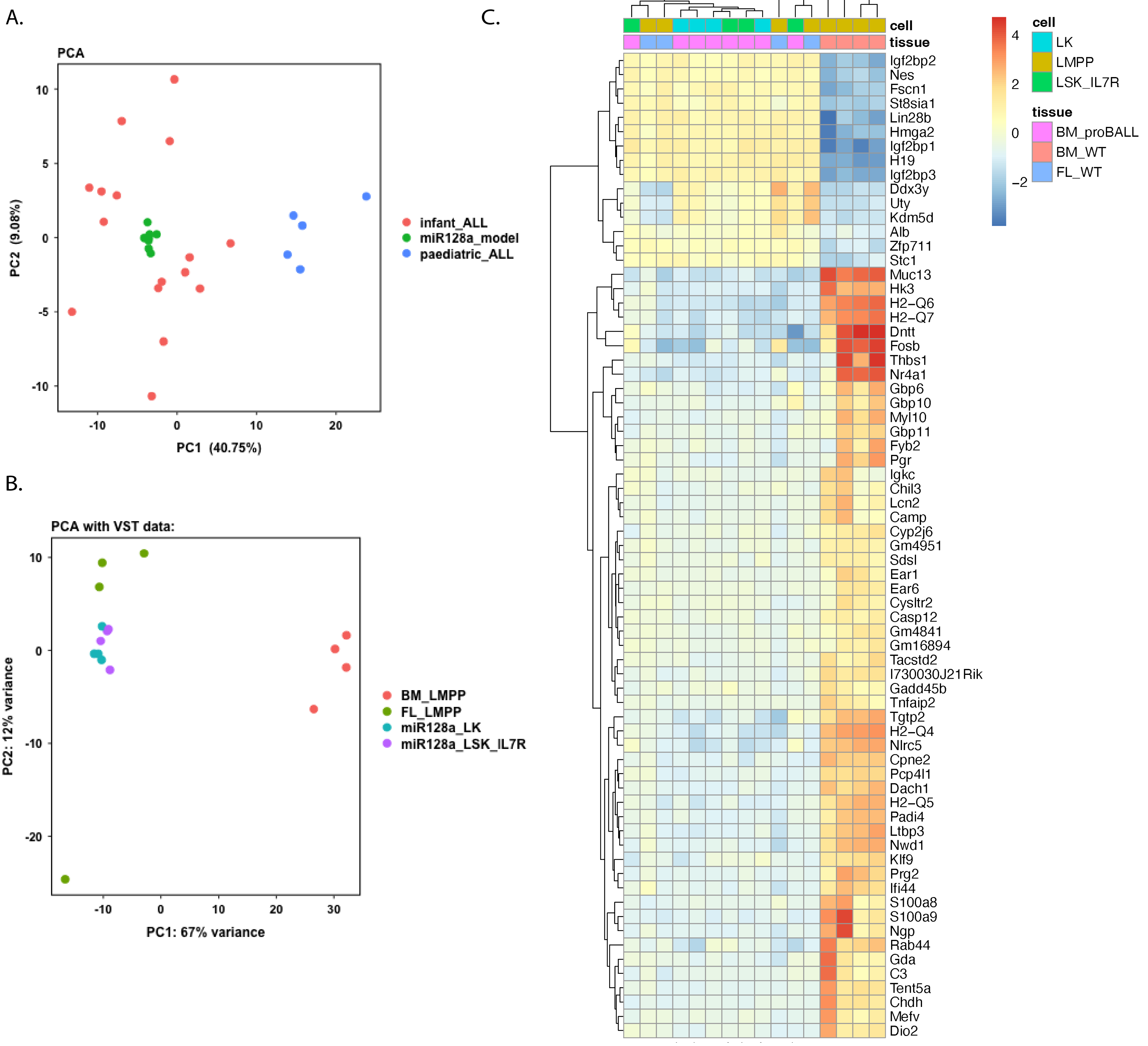

Supplemental Figure 4

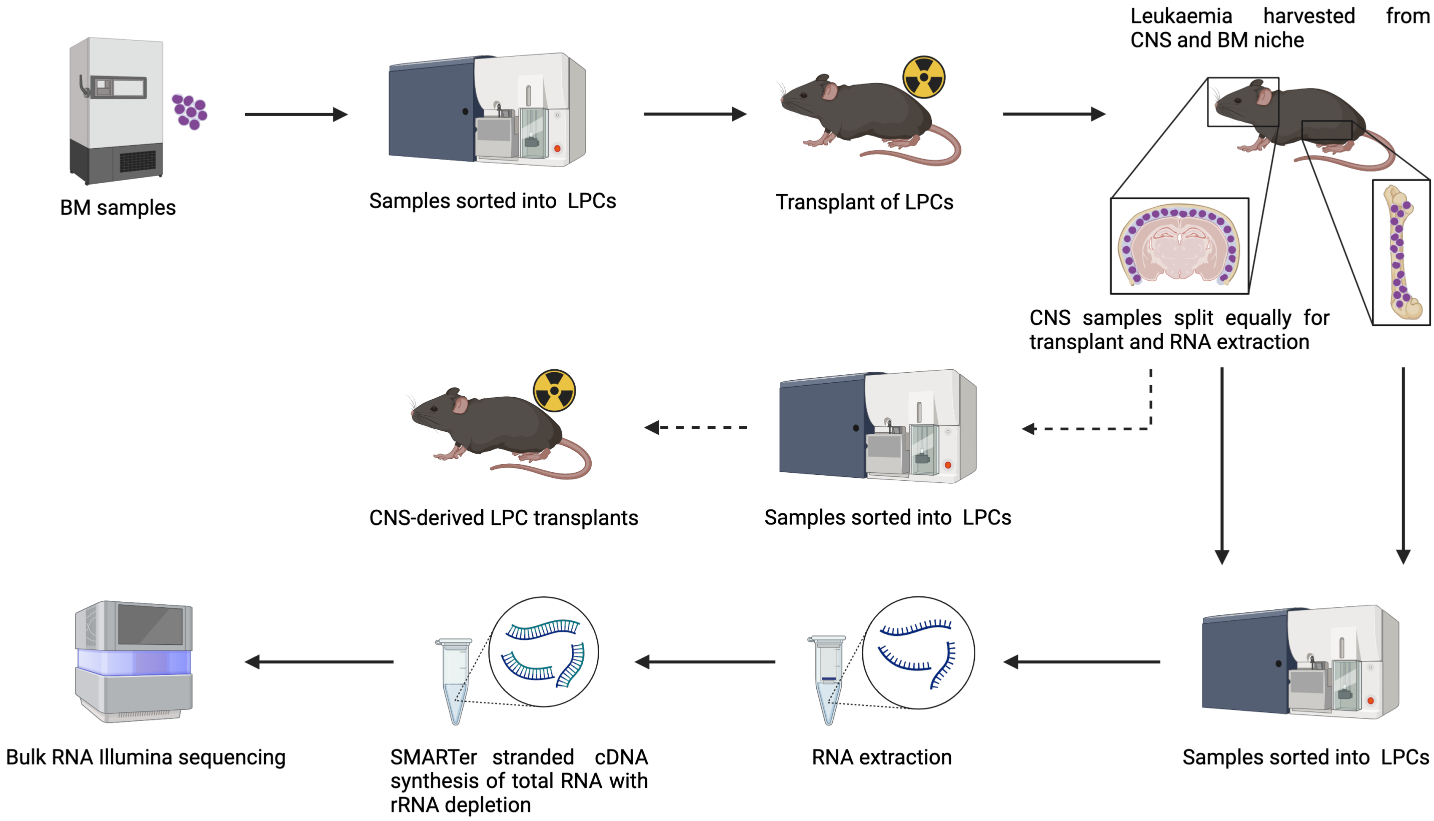


| Model | Cell type | Cell number - BM | Cell number - CNS |
|---|---|---|---|
| miR-128a | LSK IL7R | 800,000 | 135,000-800,000 |
| miR-128a | LK/CLP | 800,000 | 224,000-800,000 |

Supplemental Figure 5

| miRNA family | p-value | FDR |
|---|---|---|
| mmu-miR-296-3p | 3.41151E-05 | 0.003808817 |
| mmu-miR-326-3p/mmu-miR-330-5p | 5.34683E-05 | 0.003808817 |
| mmu-miR-1957b/mmu-miR-1a-3p/mmu-miR-206-3p/mmu-miR-6349/mmu-miR-6382 | 7.57974E-05 | 0.003808817 |
| mmu-miR-374b-5p | 5.76735E-05 | 0.003808817 |
| mmu-miR-212-5p | 0.000330344 | 0.013279839 |
| mmu-miR-10a-5p/mmu-miR-10b-5p | 0.000490045 | 0.014342643 |
| mmu-miR-134-5p | 0.000499495 | 0.014342643 |
| mmu-miR-455-3p.2/mmu-miR-682 | 0.000647999 | 0.016280969 |
| mmu-miR-101a-3p.1 | 0.000956916 | 0.021371115 |
| mmu-miR-129-1-3p/mmu-miR-129-2-3p | 0.001681479 | 0.029718622 |
| mmu-miR-25-3p/mmu-miR-32-5p/mmu-miR-363-3p/mmu-miR-367-3p/mmu-miR-92a-3p/mmu-miR-92b-3p | 0.001666795 | 0.029718622 |
| mmu-miR-340-5p | 0.001774246 | 0.029718622 |
| mmu-miR-29a-3p/mmu-miR-29b-3p/mmu-miR-29c-3p | 0.002364952 | 0.036565797 |
| mmu-miR-132-3p/mmu-miR-212-3p | 0.003032119 | 0.043532572 |
| mmu-miR-193a-3p/mmu-miR-193b-3p | 0.003501593 | 0.043988759 |
| mmu-miR-381-3p/mmu-miR-539-3p | 0.003354227 | 0.043988759 |
| mmu-miR-22-3p | 0.004337521 | 0.051284811 |
| mmu-miR-100-5p/mmu-miR-99a-5p/mmu-miR-99b-5p | 0.004653236 | 0.051961139 |
| mmu-miR-219a-2-3p | 0.006093059 | 0.064458155 |
| mmu-miR-137-3p | 0.006844861 | 0.068790851 |
| mmu-miR-331-3p | 0.007629126 | 0.073021638 |
| mmu-miR-183-5p | 0.009497906 | 0.086776326 |
| mmu-miR-101a-3p.2/mmu-miR-101b-3p.1/mmu-miR-101b-3p.2 | 0.010033682 | 0.087685653 |
| mmu-miR-183-5p.2 | 0.012477091 | 0.104495639 |
| mmu-miR-330-3p.1/mmu-miR-6240 | 0.013662785 | 0.109848794 |
| mmu-miR-18a-5p/mmu-miR-18b-5p | 0.015423751 | 0.11923746 |

| | | |
|---|---|---|
| mmu-miR-130a-3p/mmu-miR-130b-3p/mmu-miR-130c/mmu-miR-301a-3p/mmu-miR-301b-3p/mmu-miR-6341/mmu-miR-6389/mmu-miR-721 | 0.016490875 | 0.1227654 |
| mmu-miR-106a-5p/mmu-miR-106b-5p/mmu-miR-17-5p/mmu-miR-20a-5p/mmu-miR-20b-5p/mmu-miR-6383/mmu-miR-93-5p | 0.019646319 | 0.131443716 |
| mmu-miR-187-3p | 0.020247305 | 0.131443716 |
| mmu-miR-145a-5p/mmu-miR-145b | 0.019195744 | 0.131443716 |
| mmu-miR-488-3p | 0.02038538 | 0.131443716 |
| mmu-miR-129-5p | 0.020926363 | 0.131443716 |
| mmu-miR-382-3p | 0.023219218 | 0.134725256 |
| mmu-miR-423-5p/mmu-miR-6921-5p | 0.022707293 | 0.134725256 |
| mmu-miR-153-3p | 0.023459622 | 0.134725256 |
| mmu-miR-34a-5p/mmu-miR-34b-5p/mmu-miR-34c-5p/mmu-miR-449a-5p/mmu-miR-449b/mmu-miR-449c-5p | 0.026708955 | 0.149124997 |
| mmu-miR-144-3p | 0.028323357 | 0.150260917 |
| mmu-miR-135a-5p/mmu-miR-135b-5p | 0.028407537 | 0.150260917 |
| mmu-miR-30a-5p/mmu-miR-30b-5p/mmu-miR-30c-5p/mmu-miR-30d-5p/mmu-miR-30e-5p/mmu-miR-384-5p | 0.029453394 | 0.151798262 |
| mmu-miR-124-3p.2/mmu-miR-5624-3p/mmu-miR-6540-5p | 0.030480308 | 0.153163549 |
| mmu-miR-290a-5p/mmu-miR-292a-5p/mmu-miR-293-5p | 0.0332196 | 0.162857064 |
| mmu-miR-346-5p | 0.036729941 | 0.175779004 |
| mmu-miR-486a-5p/mmu-miR-486b-5p | 0.04241543 | 0.198267477 |
| mmu-miR-490-3p | 0.048173015 | 0.219617106 |

Supplemental Figure 6

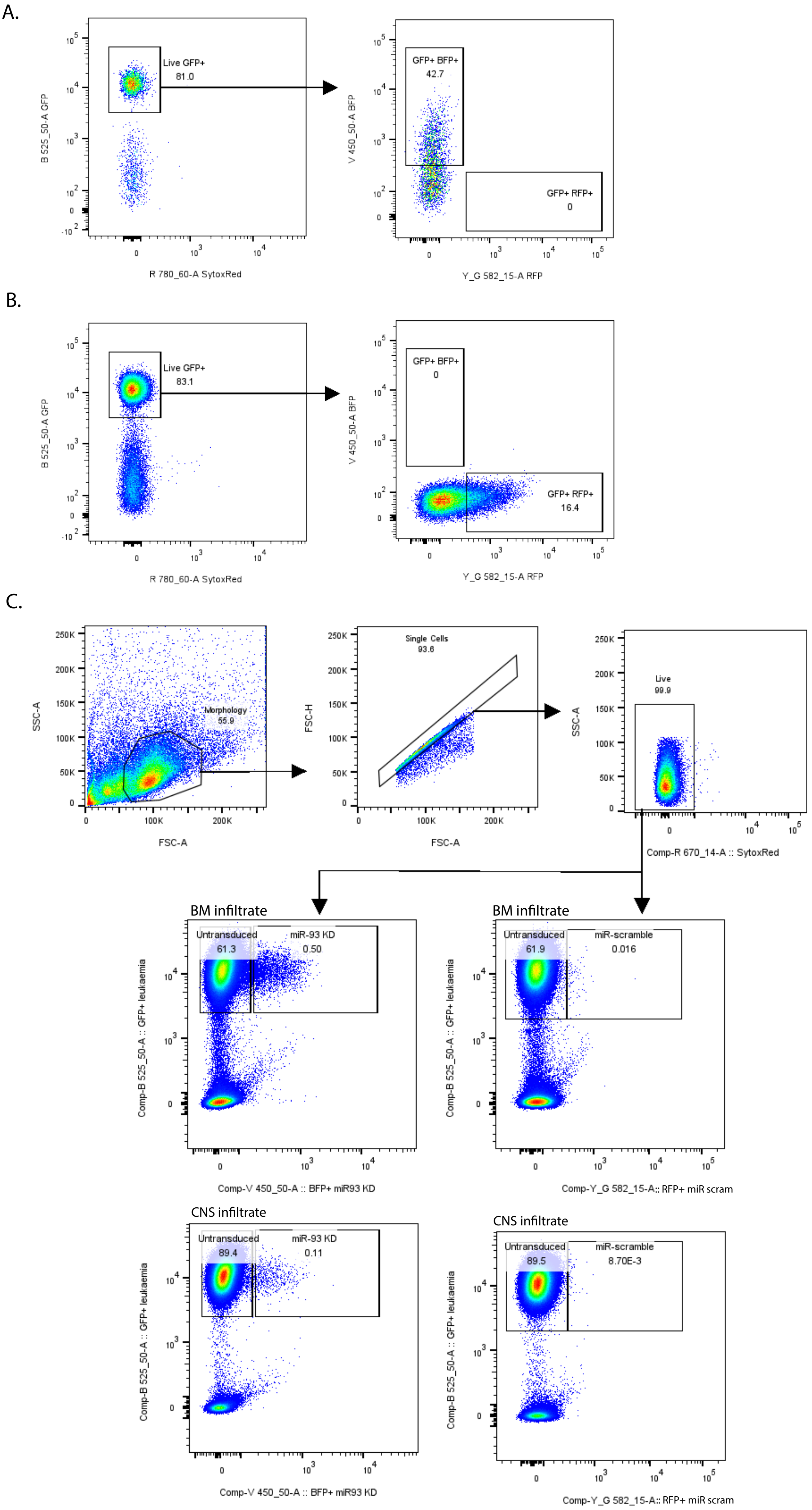

**Supplemental Table 1. Transplant details of mice used for RNA sequencing and LPC comparisons.**

| identifier | age at transplant | sex | date of transplant | source of cells | LPC type | number of cells transplanted | origin of cells | recipient number | date of cull | type of leuakemia | terminal leukaemia development? | BM + CNS RNA sequencing |
|---|---|---|---|---|---|---|---|---|---|---|---|---|
| AD1.1 EM1 | 9 weeks 5 days | F | 21.10.20 | BM | LK/CLP | 25 000 | CM58.2 EM13 | quaternary | 27.11.20 | miR-128a Kmt2a-AFF1+ | Y | N |
| AD1.1 EM3 | 9 weeks 5 days | F | 21.10.20 | BM | LK/CLP | 25 000 | CM58.2 EM13 | quaternary | 27.11.20 | miR-128a Kmt2a-AFF1+ | Y | N |
| AD1.1 EM4 | 9 weeks 5 days | F | 21.10.20 | BM | LK/CLP | 25 000 | CM58.2 EM13 | quaternary | 20.11.20 | miR-128a Kmt2a-AFF1+ | Y | N |
| AD1.1 EM10 | 9 weeks 5 days | F | 21.10.20 | BM | LK/CLP | 25 000 | CM58.2 EM13 | quaternary | 18.11.20 | miR-128a Kmt2a-AFF1+ | Y | N |
| AD1.1 EM11 | 9 weeks 5 days | F | 21.10.20 | BM | LK/CLP | 25 000 | CM58.2 EM13 | quaternary | 19.11.20 | miR-128a Kmt2a-AFF1+ | Y | N |
| AD1.1 EM13 | 9 weeks 5 days | F | 21.10.20 | BM | LK/CLP | 25 000 | CM58.2 EM13 | quaternary | 27.11.20 | miR-128a Kmt2a-AFF1+ | Y | N |
| AD1.2 EM31 | 9 weeks 5 days | F | 21.10.20 | BM | LK/CLP | 25 000 | CM58.2 EM13 | quaternary | 25.11.20 | miR-128a Kmt2a-AFF1+ | Y | Y |
| AD1.2 EM1 | 9 weeks 5 days | F | 21.10.20 | BM | LSK IL7R | 10 000 | CM58.2 EM13 | quaternary | 25.11.20 | miR-128a Kmt2a-AFF1+ | Y | Y |
| AD1.2 EM3 | 9 weeks 5 days | F | 21.10.20 | BM | LSK IL7R | 10 000 | CM58.2 EM13 | quaternary | 18.11.20 | miR-128a Kmt2a-AFF1+ | Y | N |
| AD1.2 EM4 | 9 weeks 5 days | F | 21.10.20 | BM | LSK IL7R | 1 700 | CM58.2 EM13 | quaternary | 19.11.20 | miR-128a Kmt2a-AFF1+ | Y | N |
| AD1.2 EM11 | 9 weeks 5 days | F | 21.10.20 | BM | LSK IL7R | 10 000 | CM58.2 EM13 | quaternary | 27.11.20 | miR-128a Kmt2a-AFF1+ | Y | N |
| AD1.2 EM14 | 9 weeks 5 days | F | 21.10.20 | BM | LSK IL7R | 10 000 | CM58.2 EM13 | quaternary | 27.11.20 | miR-128a Kmt2a-AFF1+ | Y | N |
| AD2.1 EM14 | 11 weeks 1 day | F | 17.11.20 | BM | LSK IL7R | 17 500 | CM56 EM4 | quaternary | 14.12.20 | miR-128a Kmt2a-AFF1+ | Y | N |
| AD2.1 EM30 | 11 weeks 0 days | F | 17.11.20 | BM | LSK IL7R | 17 500 | CM56 EM4 | quaternary | 14.12.20 | miR-128a Kmt2a-AFF1+ | Y | N |
| AD2.1 EM31 | 11 weeks 0 days | F | 17.11.20 | BM | LSK IL7R | 17 500 | CM56 EM4 | quaternary | 14.12.20 | miR-128a Kmt2a-AFF1+ | Y | Y |
| AD2.1 EM34 | 11 weeks 2 days | F | 17.11.20 | BM | LSK IL7R | 17 500 | CM56 EM4 | quaternary | 14.12.20 | miR-128a Kmt2a-AFF1+ | Y | N |
| AD2.1 EM41 | 11 weeks 2 days | F | 17.11.20 | BM | LK/CLP | 52 000 | CM56 EM4 | quaternary | 16.12.20 | miR-128a Kmt2a-AFF1+ | Y | Y |
| AD2.1 EM44 | 11 weeks 2 days | F | 17.11.20 | BM | LK/CLP | 52 000 | CM56 EM4 | quaternary | 14.12.20 | miR-128a Kmt2a-AFF1+ | Y | N |
| AD2.2 EM1 | 12 weeks 0 days | F | 17.11.20 | BM | LK/CLP | 52 000 | CM56 EM4 | quaternary | 23.12.20 | miR-128a Kmt2a-AFF1+ | Y | N |
| AD2.2 EM3 | 12 weeks 0 days | F | 17.11.20 | BM | LK/CLP | 52 000 | CM56 EM4 | quaternary | 23.12.20 | miR-128a Kmt2a-AFF1+ | Y | N |
| AD4 EM0 | 6 weeks 5 days | F | 13.1.21 | BM | LSK IL7R | 150 000 | CM56 EM3 | quaternary | 8.2.21 | miR-128a Kmt2a-AFF1+ | Y | Y |
| AD4 EM1 | 6 weeks 5 days | F | 13.1.21 | BM | LK/CLP | 48 000 | CM56 EM3 | quaternary | 7.2.21 | miR-128a Kmt2a-AFF1+ | Y | N |
| AD4 EM4 | 6 weeks 5 days | F | 13.1.21 | BM | LSK IL7R | 150 000 | CM56 EM3 | quaternary | 8.2.21 | miR-128a Kmt2a-AFF1+ | Y | N |
| AD4 EM11 | 6 weeks 5 days | F | 13.1.21 | BM | LK/CLP | 48 000 | CM56 EM3 | quaternary | 8.2.21 | miR-128a Kmt2a-AFF1+ | Y | Y |
| AD5 EM1 | 11 weeks 2 days | M | 21.1.21 | BM | LSK IL7R | 80 000 | CM56 EM10 | quaternary | 11.2.21 | miR-128a Kmt2a-AFF1+ | Y | Y |
| AD5 EM3 | 10 weeks 4 days | M | 21.1.21 | BM | LK/CLP | 70 000 | CM56 EM10 | quaternary | 11.2.21 | miR-128a Kmt2a-AFF1+ | Y | Y |
| AD5 EM10 | 10 weeks 4 days | M | 21.1.21 | BM | LSK IL7R | 80 000 | CM56 EM10 | quaternary | 18.2.21 | miR-128a Kmt2a-AFF1+ | Y | N |
| AD5 EM11 | 10 weeks 4 days | M | 21.1.21 | BM | LK/CLP | 70 000 | CM56 EM10 | quaternary | 18.2.21 | miR-128a Kmt2a-AFF1+ | Y | N |
| AD11 EM1 | 10 weeks 4 days | F | 18.6.21 | CNS | LSK IL7R | 45 000 | AD1.2 EM1 | quinary | 19.7.21 | miR-128a Kmt2a-AFF1+ | Y | N |
| AD11 EM3 | 10 weeks 4 days | F | 18.6.21 | CNS | LK/CLP | 51 000 | AD1.2 EM31 | quinary | 19.7.21 | miR-128a Kmt2a-AFF1+ | Y | N |
| AD11 EM4 | 10 weeks 4 days | F | 18.6.21 | CNS | LSK IL7R | 40 500 | AD1.2 EM31 | quinary | 15.7.21 | miR-128a Kmt2a-AFF1+ | Y | N |
| AD11 EM10 | 10 weeks 4 days | F | 18.6.21 | CNS | LK/CLP | 51 000 | AD1.2 EM31 | quinary | 17.7.21 | miR-128a Kmt2a-AFF1+ | Y | N |
| AD11 EM11 | 10 weeks 4 days | F | 18.6.21 | CNS | LSK IL7R | 45 000 | AD1.2 EM1 | quinary | 19.7.21 | miR-128a Kmt2a-AFF1+ | Y | N |
| AD11 EM13 | 10 weeks 4 days | F | 18.6.21 | CNS | LSK IL7R | 40 500 | AD1.2 EM31 | quinary | 15.7.21 | miR-128a Kmt2a-AFF1+ | Y | N |
| AD12.3 EM0 | 9 weeks 6 days | M | 24.6.21 | CNS | LSK IL7R | 50 000 | AD5 EM1 | quinary | 26.7.21 | miR-128a Kmt2a-AFF1+ | Y | N |
| AD12.3 EM1 | 9 weeks 6 days | M | 24.6.21 | CNS | LK/CLP | 70 000 | AD5 EM3 | quinary | 26.7.21 | miR-128a Kmt2a-AFF1+ | Y | N |
| AD12.3 EM3 | 9 weeks 6 days | M | 24.6.21 | CNS | LSK IL7R | 50 000 | AD5 EM1 | quinary | 26.7.21 | miR-128a Kmt2a-AFF1+ | Y | N |
| AD12.3 EM4 | 9 weeks 6 days | M | 24.6.21 | CNS | LK/CLP | 70 000 | AD5 EM3 | quinary | 26.7.21 | miR-128a Kmt2a-AFF1+ | Y | N |
| AD14 EM1 | 11 week 4 days | F | 7.7.21 | CNS | LK/CLP | 38 000 | AD4 EM11 | quinary | 4.8.21 | miR-128a Kmt2a-AFF1+ | Y | N |
| AD14 EM3 | 11 week 4 days | F | 7.7.21 | CNS | LSK IL7R | 180 000 | AD4 NM | quinary | 2.8.21 | miR-128a Kmt2a-AFF1+ | Y | N |
| AD14 EM4 | 11 week 4 days | F | 7.7.21 | CNS | LK/CLP | 20 000 | AD2.1 EM41 | quinary | 2.8.21 | miR-128a Kmt2a-AFF1+ | Y | N |
| AD14 EM10 | 11 week 4 days | F | 7.7.21 | CNS | LK/CLP | 20 000 | AD2.1 EM41 | quinary | 2.8.21 | miR-128a Kmt2a-AFF1+ | Y | N |
| AD14 EM11 | 10 weeks 2 days | F | 7.7.21 | CNS | LK/CLP | 38 000 | AD4 EM11 | quinary | 31.7.21 | miR-128a Kmt2a-AFF1+ | Y | N |
| AD14 EM13 | 10 weeks 2 days | F | 7.7.21 | CNS | LSK IL7R | 180 000 | AD4 NM | quinary | 2.8.21 | miR-128a Kmt2a-AFF1+ | Y | N |

**Supplemental Table 2. Differentially expressed genes between BM-derived LSK IL7R and LK/CLP cells.**

| geneid | baseMean | log2FoldChange | lfcSE | stat | pvalue | padj | symbol | genename |
|---|---|---|---|---|---|---|---|---|
| ENSMUSG00000026605 | 4580 121679 | 0 689363978 | 0 109861344 | 6 274854743 | 3 50E-10 | 5 06E-06 | Cenpf | centromere protein F |
| ENSMUSG00000018362 | 1866 57347 | 0 578843046 | 0 097999775 | 5 906575288 | 3 49E-09 | 2 52E-05 | Kpna2 | karyopherin (importin) alpha 2 |
| ENSMUSG00000041431 | 792 5394675 | 0 629856787 | 0 115825228 | 5 437993062 | 5.38840722027565e-08 | 0 000259524 | Ccnb1 | cyclin B1 |
| ENSMUSG00000030867 | 810 8829769 | 0 651256633 | 0 123739131 | 5 263142129 | 1.41614016276501e-07 | 0 000511545 | Plk1 | polo like kinase 1 |
| ENSMUSG00000001403 | 732 5902558 | 0 714610897 | 0 136946128 | 5 218189859 | 1.80680207952816e-07 | 0 00052213 | Ube2c | ubiquitin-conjugating enzyme E2C |
| ENSMUSG00000033952 | 2207 374747 | 0 534225737 | 0 103184528 | 5 177382169 | 2.25020971607362e-07 | 0 000541888 | Aspm | abnormal spindle microtubule assembly |
| ENSMUSG00000066878 | 967 6170373 | 0 582787718 | 0 114599511 | 5 085429382 | 3.66795291213403e-07 | 0 000757118 | Gm10184 | karyopherin (importin) alpha 2 pseudogene |
| ENSMUSG00000019942 | 978 525135 | 0 595132182 | 0 121546661 | 4 896326875 | 9.76445937075272e-07 | 0 00148944 | Cdk1 | cyclin-dependent kinase 1 |
| ENSMUSG00000069083 | 827 5275081 | 0 563660281 | 0 115370326 | 4 885660797 | 1.03082536653458e-06 | 0 00148944 | Gm10259 | karyopherin (importin) alpha 2 pseudogene |
| ENSMUSG00000037725 | 453 2323801 | 0 567499334 | 0 119077445 | 4 765800376 | 1.88105429401257e-06 | 0 00247085 | Ckap2 | cytoskeleton associated protein 2 |
| ENSMUSG00000026683 | 919 7506723 | 0 502311718 | 0 106733657 | 4 706216687 | 2.52356282209251e-06 | 0 002804843 | Nuf2 | NUF2, NDC80 kinetochore complex component |
| ENSMUSG00000051378 | 780 1817219 | 0 540119292 | 0 114684389 | 4 709614756 | 2.48185447258163e-06 | 0 002804843 | Kif18b | kinesin family member 18B |
| ENSMUSG00000027469 | 2366 981345 | 0 512661567 | 0 112859371 | 4 542481151 | 5.55959690009019e-06 | 0 005737901 | Tpx2 | TPX2, microtubule-associated |
| ENSMUSG00000036752 | 2933 355166 | 0 515021135 | 0 114005829 | 4 517498259 | 6.25745423443673e-06 | 0 006027597 | Tubb4b | tubulin, beta 4B class IVB |
| ENSMUSG00000030337 | 529 5886569 | -0 482037924 | 0 108703452 | -4 434430699 | 9.23159083893208e-06 | 0 008336704 | Vamp1 | vesicle-associated membrane protein 1 |
| ENSMUSG00000031004 | 27856 56849 | 0 382492509 | 0 088165519 | 4 338345818 | 1.43559142791104e-05 | 0 011523811 | Mki67 | antigen identified by monoclonal antibody Ki 67 |
| ENSMUSG00000020330 | 1162 313821 | 0 552835304 | 0 128338081 | 4 307648205 | 1.64999536637098e-05 | 0 012547781 | Hmmr | hyaluronan mediated motility receptor (RHAMM) |
| ENSMUSG00000006398 | 679 8080701 | 0 579014486 | 0 135342413 | 4 278145123 | 1.88457144980339e-05 | 0 012565301 | Cdc20 | cell division cycle 20 |
| ENSMUSG00000027330 | 1926 048472 | 0 402415135 | 0 094038523 | 4 279258342 | 1.87517110331265e-05 | 0 012565301 | Cdc25b | cell division cycle 25B |
| ENSMUSG00000079553 | 919 7185952 | 0 529201869 | 0 123796037 | 4 274788454 | 1.91318863038467e-05 | 0 012565301 | Kifc1 | kinesin family member C1 |
| ENSMUSG00000027306 | 1340 959046 | 0 588626349 | 0 13916717 | 4 229635122 | 2.34070652718127e-05 | 0 014704725 | Nusap1 | nucleolar and spindle associated protein 1 |
| ENSMUSG00000025154 | 567 9049728 | 0 451775796 | 0 107828681 | 4 189755369 | 2.79255349754647e-05 | 0 016812336 | Arhgap19 | Rho GTPase activating protein 19 |
| ENSMUSG00000024795 | 1397 099882 | 0 408353892 | 0 09791792 | 4 170369328 | 3.04106389522841e-05 | 0 016900128 | Kif20b | kinesin family member 20B |
| ENSMUSG00000045328 | 2475 801999 | 0 381910744 | 0 091432872 | 4 176952297 | 2.9544095366027e-05 | 0 016900128 | Cenpe | centromere protein E |
| ENSMUSG00000024660 | 3969 410429 | 0 414189876 | 0 099780394 | 4 151014645 | 3.31004583101916e-05 | 0 017713649 | Incenp | inner centromere protein |
| ENSMUSG00000029177 | 813 0701371 | 0 458774964 | 0 11082676 | 4 139568491 | 3.47959705637379e-05 | 0 017955964 | Cenpa | centromere protein A |
| ENSMUSG00000021697 | 266 7090784 | 0 600715635 | 0 146654074 | 4 096140105 | 4.20095747568069e-05 | 0 018334515 | Depdc1b | DEP domain containing 1B |
| ENSMUSG00000026622 | 683 0214053 | 0 534757 | 0 130963861 | 4 08324094 | 4.44119340223971e-05 | 0 018334515 | Nek2 | NIMA (never in mitosis gene a)-related expressed kinase 2 |
| ENSMUSG00000027326 | 2130 601023 | 0 354183396 | 0 086678687 | 4 086164741 | 4.38562439302831e-05 | 0 018334515 | Knl1 | kinetochore scaffold 1 |
| ENSMUSG00000030654 | 1221 830477 | 0 475112118 | 0 115493775 | 4 113746531 | 3.89288873065843e-05 | 0 018334515 | Arl6ip1 | ADP-ribosylation factor-like 6 interacting protein 1 |
| ENSMUSG00000041498 | 998 9975823 | 0 54368994 | 0 132995345 | 4 088037368 | 4.35038088904573e-05 | 0 018334515 | Kif14 | kinesin family member 14 |
| ENSMUSG00000069892 | 128 9961604 | 0 702872499 | 0 171203648 | 4 105476175 | 4.03482869478102e-05 | 0 018334515 | 9930111J21R | RIKEN cDNA 9930111J21 gene 2 |
| ENSMUSG00000074802 | 360 1594657 | 0 498458823 | 0 121080646 | 4 116750618 | 3.84251457013772e-05 | 0 018334515 | Gas2l3 | growth arrest-specific 2 like 3 |
| ENSMUSG00000032218 | 871 7818571 | 0 440284813 | 0 10905823 | 4 037153468 | 5.41036776293983e-05 | 0 021128217 | Ccnb2 | cyclin B2 |
| ENSMUSG00000038943 | 1189 187038 | 0 438141201 | 0 108526602 | 4 037177913 | 5.40980424961033e-05 | 0 021128217 | Prc1 | protein regulator of cytokinesis 1 |
| ENSMUSG00000119084 | 98 12731932 | -0 795284047 | 0 197898039 | -4 01865551 | 5.85311730214556e-05 | 0 022255708 | Gm26317 | predicted gene, 26317 |
| ENSMUSG00000022385 | 389 1872864 | 0 580939248 | 0 145290104 | 3 998477751 | 6.3751172334313e-05 | 0 023322828 | Gtse1 | G two S phase expressed protein 1 |
| ENSMUSG00000034311 | 1051 541483 | 0 468710897 | 0 117310567 | 3 99547038 | 6.45659295389789e-05 | 0 023322828 | Kif4 | kinesin family member 4 |
| ENSMUSG00000041219 | 2253 776003 | 0 363571889 | 0 092453121 | 3 9324999 | 8.40670147716821e-05 | 0 029626446 | Arhgap11a | Rho GTPase activating protein 11A |
| ENSMUSG00000020808 | 393 2741764 | 0 594938697 | 0 151599638 | 3 924407113 | 8.69435975131009e-05 | 0 029910668 | Pimreg | PICALM interacting mitotic regulator |
| ENSMUSG00000012443 | 3315 564116 | 0 413883609 | 0 105819948 | 3 911205916 | 9.18364312568508e-05 | 0 030859177 | Kif11 | kinesin family member 11 |
| ENSMUSG00000118723 | 131 163208 | -0 712455213 | 0 183193847 | -3 889078295 | 0 000100626 | 0 033044093 | Gm25127 | predicted gene, 25127 |
| ENSMUSG00000027496 | 622 6816485 | 0 513357248 | 0 13286748 | 3 863678665 | 0 000111692 | 0 035863102 | Aurka | aurora kinase A |
| ENSMUSG00000020914 | 14842 43907 | 0 425901285 | 0 110792516 | 3 844134072 | 0 000120979 | 0 038000511 | Top2a | topoisomerase (DNA) II alpha |
| ENSMUSG00000036964 | 66 26790713 | -0 961172596 | 0 250439582 | -3 837942028 | 0 00012407 | 0 038142213 | Trim17 | tripartite motif-containing 17 |
| ENSMUSG00000027883 | 440 9884041 | 0 440447712 | 0 11504941 | 3 82833524 | 0 000129013 | 0 038835586 | Gpsm2 | G-protein signalling modulator 2 (AGS3-like, C. elegans) |
| ENSMUSG00000034573 | 789 6676632 | -0 648271103 | 0 169698747 | -3 820128981 | 0 000133382 | 0 03933133 | Ptpn13 | protein tyrosine phosphatase, non-receptor type 13 |
| ENSMUSG00000028175 | 335 9847201 | 0 488097209 | 0 128843194 | 3 788304172 | 0 000151679 | 0 04383222 | Depdc1a | DEP domain containing 1a |
| ENSMUSG00000062248 | 540 9503835 | 0 438808436 | 0 116320883 | 3 772396001 | 0 000161687 | 0 04580825 | Cks2 | CDC28 protein kinase regulatory subunit 2 |
| ENSMUSG00000037544 | 784 9267428 | 0 433668537 | 0 115147654 | 3 766195162 | 0 000165754 | 0 046057367 | Dlgap5 | DLG associated protein 5 |
| ENSMUSG00000020493 | 297 4234106 | 0 50334808 | 0 134128406 | 3 752732888 | 0 000174917 | 0 047240473 | Prr11 | proline rich 11 |
| ENSMUSG00000020737 | 1284 545506 | 0 380997583 | 0 10215049 | 3 729767571 | 0 000191656 | 0 049450795 | Jpt1 | Jupiter microtubule associated homolog 1 |

**Supplemental Table 3. Differentially expressed genes between CNS-derived LSK IL7R and LK/CLP cells.**

| geneid | baseMean | log2FoldChange | lfcSE | stat | pvalue | padj | symbol | genename |
|---|---|---|---|---|---|---|---|---|
| ENSMUSG00000047907 | 21 0957264 | 2 260487861 | 0 454270847 | 4 976079524 | 6 49E-07 | 0 017718147 | Tshz2 | teashirt zinc finger family member 2 |
| ENSMUSG00000001403 | 787 6874 | 0 654997575 | 0 140397251 | 4 665316232 | 3 08E-06 | 0 030374436 | Ube2c | ubiquitin-conjugating enzyme E2C |
| ENSMUSG00000033278 | 20 9795858 | 2 005284419 | 0 436956608 | 4 589207214 | 4 45E-06 | 0 030374436 | Ptprm | protein tyrosine phosphatase, receptor type, M |
| ENSMUSG00000037974 | 73 3962314 | 1 474888696 | 0 320109247 | 4 607454206 | 4 08E-06 | 0 030374436 | Muc5ac | mucin 5, subtypes A and C, tracheobronchial/gastric |

**Supplemental Table 4. Differentially expressed genes between CNS-derived and BM-derived LPCs.**

| geneid | baseMean | log2FoldChange | lfcSE | stat | pvalue | padj | symbol | genename |
|---|---|---|---|---|---|---|---|---|
| ENSMUSG00000026728 | 3266 262593 | -0 671097124 | 0 06262116 | -10 71677892 | 8 49E-27 | 1 71E-22 | Vim | vimentin |
| ENSMUSG00000033066 | 823 9312614 | -0 713484152 | 0 082087428 | -8 691759165 | 3 57E-18 | 3 60E-14 | Gas7 | growth arrest specific 7 |
| ENSMUSG00000026600 | 544 7933161 | 0 593538078 | 0 073059664 | 8 124018776 | 4 51E-16 | 3 04E-12 | Soat1 | sterol O-acyltransferase 1 |
| ENSMUSG00000024900 | 3342 977048 | -0 452030445 | 0 057100221 | -7 91643953 | 2 44E-15 | 1 05E-11 | Cpt1a | carnitine palmitoyltransferase 1a, liver |
| ENSMUSG00000025014 | 1823 445445 | 0 958323104 | 0 121167259 | 7 909092884 | 2 59E-15 | 1 05E-11 | Dntt | deoxynucleotidyltransferase, terminal |
| ENSMUSG00000027985 | 4177 033657 | 0 314120775 | 0 040819955 | 7 695274841 | 1 41E-14 | 4 75E-11 | Lef1 | lymphoid enhancer binding factor 1 |
| ENSMUSG00000038587 | 1131 019673 | -0 423005607 | 0 055727076 | -7 590665756 | 3 18E-14 | 9 18E-11 | Akap12 | A kinase (PRKA) anchor protein (gravin) 12 |
| ENSMUSG00000002885 | 911 627176 | -0 462313973 | 0 061084433 | -7 568441769 | 3 78E-14 | 9 53E-11 | Adgre5 | adhesion G protein-coupled receptor E5 |
| ENSMUSG00000054364 | 80 1795637 | -1 52989375 | 0 203584196 | -7 514796219 | 5 70E-14 | 1 28E-10 | Rhob | ras homolog family member B |
| ENSMUSG00000000915 | 1903 08103 | -0 335011574 | 0 046227493 | -7 247020118 | 4 26E-13 | 8 12E-10 | Hip1r | huntingtin interacting protein 1 related |
| ENSMUSG00000027858 | 1002 51043 | -0 425152929 | 0 058707625 | -7 241868981 | 4 43E-13 | 8 12E-10 | Tspan2 | tetraspanin 2 |
| ENSMUSG00000026011 | 16 34873733 | 3 889101642 | 0 549669587 | 7 075344411 | 1 49E-12 | 2 51E-09 | Ctla4 | cytotoxic T-lymphocyte-associated protein 4 |
| ENSMUSG00000069833 | 31964 07344 | -0 303907373 | 0 043583785 | -6 972945928 | 3 10E-12 | 4 82E-09 | Ahnak | AHNAK nucleoprotein (desmoyokin) |
| ENSMUSG00000016756 | 1247 299252 | -0 345879366 | 0 050598083 | -6 835819612 | 8 15E-12 | 1 18E-08 | Cmah | cytidine monophospho-N-acetylneuraminic acid hydroxylase |
| ENSMUSG00000001281 | 528 587906 | -0 580031218 | 0 085983244 | -6 74586335 | 1 52E-11 | 2 00E-08 | Itgb7 | integrin beta 7 |
| ENSMUSG00000030616 | 2694 534136 | -0 347426573 | 0 05154758 | -6 739920148 | 1 58E-11 | 2 00E-08 | Sytl2 | synaptotagmin-like 2 |
| ENSMUSG00000078942 | 1842 425579 | -0 40967692 | 0 063576996 | -6 44379177 | 1 17E-10 | 1 38E-07 | Naip6 | NLR family, apoptosis inhibitory protein 6 |
| ENSMUSG00000082956 | 476 870824 | -0 491461276 | 0 076438597 | -6 429491073 | 1 28E-10 | 1 44E-07 | NA | NA |
| ENSMUSG00000002489 | 1109 333482 | 0 303250645 | 0 047517024 | 6 38193681 | 1 75E-10 | 1 86E-07 | Tiam1 | T cell lymphoma invasion and metastasis 1 |
| ENSMUSG00000025905 | 659 9169454 | -0 471843111 | 0 075100868 | -6 282791686 | 3 33E-10 | 3 36E-07 | Oprk1 | opioid receptor, kappa 1 |
| ENSMUSG00000030523 | 323 0818538 | -0 526577754 | 0 086183622 | -6 109951559 | 9 97E-10 | 8 97E-07 | Trpm1 | transient receptor potential cation channel, subfamily M, member 1 |
| ENSMUSG00000031328 | 7180 682866 | -0 257394573 | 0 042060326 | -6 119652389 | 9 38E-10 | 8 97E-07 | Flna | filamin, alpha |
| ENSMUSG00000059146 | 432 4487716 | -0 496653463 | 0 081339219 | -6 105953168 | 1 02E-09 | 8 97E-07 | Ntrk3 | neurotrophic tyrosine kinase, receptor, type 3 |
| ENSMUSG00000017639 | 379 8778067 | 0 618318339 | 0 102222835 | 6 048730117 | 1 46E-09 | 1 23E-06 | Rab11fip4 | RAB11 family interacting protein 4 (class II) |
| ENSMUSG00000024030 | 1016 119974 | 0 464271504 | 0 079467606 | 5 842273718 | 5 15E-09 | 4 16E-06 | Abcg1 | ATP binding cassette subfamily G member 1 |
| ENSMUSG00000075254 | 3734 794258 | 0 215839619 | 0 037365834 | 5 776389733 | 7 63E-09 | 5 93E-06 | Heg1 | heart development protein with EGF-like domains 1 |
| ENSMUSG00000005087 | 1167 467086 | -0 278358865 | 0 048964452 | -5 684917347 | 1 31E-08 | 9 79E-06 | Cd44 | CD44 antigen |
| ENSMUSG00000035711 | 582 5347634 | -0 427215424 | 0 078444913 | -5 44605646 | 5 15E-08 | 3 71E-05 | Dok3 | docking protein 3 |
| ENSMUSG00000020601 | 305 7523441 | 0 587193939 | 0 108055371 | 5 434194836 | 5 50E-08 | 3 83E-05 | Trib2 | tribbles pseudokinase 2 |
| ENSMUSG00000054150 | 413 8799115 | -0 386781687 | 0 071450334 | -5 413294292 | 6 19E-08 | 4 16E-05 | Syne3 | spectrin repeat containing, nuclear envelope family member 3 |
| ENSMUSG00000024431 | 2602 710078 | -0 23486934 | 0 043541088 | -5 394200065 | 6 88E-08 | 4 48E-05 | Nr3c1 | nuclear receptor subfamily 3, group C, member 1 |
| ENSMUSG00000036523 | 83 46144851 | 0 83161071 | 0 154906496 | 5 368468917 | 7 94E-08 | 4 86E-05 | Greb1 | gene regulated by estrogen in breast cancer protein |
| ENSMUSG00000102439 | 47 87657169 | 1 269885861 | 0 236520145 | 5 369038908 | 7 92E-08 | 4 86E-05 | Flg | filaggrin |
| ENSMUSG00000074570 | 689 0027666 | -0 384588949 | 0 072302913 | -5 319134897 | 1 04E-07 | 6 19E-05 | Cass4 | Cas scaffolding protein family member 4 |
| ENSMUSG00000064147 | 108 0954797 | -0 799584737 | 0 150578422 | -5 310088446 | 1 10E-07 | 6 32E-05 | Rab44 | RAB44, member RAS oncogene family |
| ENSMUSG00000025366 | 1977 980187 | -0 270100374 | 0 051314278 | -5 263649455 | 1 41E-07 | 7 92E-05 | Esyt1 | extended synaptotagmin-like protein 1 |
| ENSMUSG00000043460 | 746 2699722 | -0 444418483 | 0 084751099 | -5 243807885 | 1 57E-07 | 8 58E-05 | Elfn2 | leucine rich repeat and fibronectin type III, extracellular 2 |
| ENSMUSG00000030265 | 816 8868826 | -0 332692667 | 0 063710338 | -5 221957329 | 1 77E-07 | 9 41E-05 | Kras | Kirsten rat sarcoma viral oncogene homolog |
| ENSMUSG00000112963 | 20 99733514 | 2 105000923 | 0 410126945 | 5 132559443 | 2 86E-07 | 0 000147964 | Gm6093 | predicted gene 6093 |
| ENSMUSG00000053617 | 624 1170679 | 0 300627497 | 0 059720002 | 5 033949886 | 4 80E-07 | 0 000242508 | Sh3pxd2a | SH3 and PX domains 2A |
| ENSMUSG00000065954 | 2969 862743 | -0 201329865 | 0 040106282 | -5 019908441 | 5 17E-07 | 0 000254559 | Tacc1 | transforming, acidic coiled-coil containing protein 1 |
| ENSMUSG00000054435 | 405 4438195 | 0 387834811 | 0 077380333 | 5 012059217 | 5 39E-07 | 0 000258855 | Gimap4 | GTPase, IMAP family member 4 |
| ENSMUSG00000028063 | 140 2606561 | -0 630414865 | 0 125993614 | -5 003546187 | 5 63E-07 | 0 000260166 | Lmna | lamin A |
| ENSMUSG00000029913 | 242 6561995 | 0 493611829 | 0 098680359 | 5 00212844 | 5 67E-07 | 0 000260166 | Prdm5 | PR domain containing 5 |
| ENSMUSG00000013663 | 5490 493434 | -0 222318183 | 0 044952555 | -4 945618372 | 7 59E-07 | 0 000340532 | Pten | phosphatase and tensin homolog |
| ENSMUSG00000016028 | 5293 858218 | -0 233392303 | 0 047448465 | -4 918858829 | 8 71E-07 | 0 000382056 | Celsr1 | cadherin, EGF LAG seven-pass G-type receptor 1 |
| ENSMUSG00000003032 | 395 3425702 | -0 53699161 | 0 109703845 | -4 894920577 | 9 83E-07 | 0 000422446 | Klf4 | Kruppel-like factor 4 (gut) |
| ENSMUSG00000018398 | 978 9279641 | -0 257775726 | 0 053011421 | -4 862645139 | 1 16E-06 | 0 000467688 | Septin8 | septin 8 |
| ENSMUSG00000020315 | 8244 421962 | -0 18856012 | 0 038734784 | -4 867979112 | 1 13E-06 | 0 000467688 | Sptbn1 | spectrin beta, non-erythrocytic 1 |
| ENSMUSG00000055148 | 44 77529304 | -1 024575655 | 0 210667113 | -4 863481722 | 1 15E-06 | 0 000467688 | Klf2 | Kruppel-like factor 2 (lung) |
| ENSMUSG00000060985 | 63 09994155 | 0 987904122 | 0 204000376 | 4 842658333 | 1 28E-06 | 0 000507154 | Tdrd5 | tudor domain containing 5 |
| ENSMUSG00000000078 | 173 5910931 | -0 52020489 | 0 107772991 | -4 826857671 | 1 39E-06 | 0 000538519 | Klf6 | Kruppel-like factor 6 |
| ENSMUSG00000032470 | 478 6016318 | -0 377675863 | 0 08011237 | -4 714326397 | 2 43E-06 | 0 000923786 | Mras | muscle and microspikes RAS |
| ENSMUSG00000024620 | 609 9938819 | -0 620201102 | 0 131972846 | -4 699459928 | 2 61E-06 | 0 000975242 | Pdgfrb | platelet derived growth factor receptor, beta polypeptide |
| ENSMUSG00000038393 | 1125 720232 | -0 371509381 | 0 079304818 | -4 684575178 | 2 81E-06 | 0 00102979 | Txnip | thioredoxin interacting protein |
| ENSMUSG00000032012 | 196 6418755 | 0 607680835 | 0 130551503 | 4 654721102 | 3 24E-06 | 0 00116959 | Nectin1 | nectin cell adhesion molecule 1 |
| ENSMUSG00000022657 | 110 8457962 | 0 57210601 | 0 123196679 | 4 6438428 | 3 42E-06 | 0 001190412 | Cd96 | CD96 antigen |
| ENSMUSG00000050271 | 40 55016425 | 1 159129518 | 0 249463325 | 4 646492707 | 3 38E-06 | 0 001190412 | Prag1 | PEAK1 related kinase activating pseudokinase 1 |
| ENSMUSG00000053641 | 947 3206438 | -0 286765754 | 0 061944566 | -4 62939324 | 3 67E-06 | 0 00125493 | Dennd4a | DENN/MADD domain containing 4A |
| ENSMUSG00000020653 | 36 29415658 | -1 077648684 | 0 233252675 | -4 620091427 | 3 84E-06 | 0 001290652 | Klf11 | Kruppel-like factor 11 |
| ENSMUSG00000020009 | 2190 362709 | -0 271618582 | 0 058860858 | -4 614587503 | 3 94E-06 | 0 001303603 | Ifngr1 | interferon gamma receptor 1 |
| ENSMUSG00000035863 | 2267 714794 | -0 244104074 | 0 052973673 | -4 608026229 | 4 07E-06 | 0 001323713 | Palm | paralemmin |
| ENSMUSG00000045216 | 398 9005628 | 0 390163696 | 0 084747685 | 4 603827203 | 4 15E-06 | 0 001329259 | Hs6st1 | heparan sulfate 6-O-sulfotransferase 1 |
| ENSMUSG00000096847 | 106 7567328 | 0 810553354 | 0 176922944 | 4 581391966 | 4 62E-06 | 0 00145705 | Tmem151b | transmembrane protein 151B |
| ENSMUSG00000035354 | 674 939717 | -0 270970413 | 0 059239261 | -4 57416933 | 4 78E-06 | 0 001480556 | Uvrag | UV radiation resistance associated gene |
| ENSMUSG00000055447 | 3769 693374 | -0 188735464 | 0 041284316 | -4 571601999 | 4 84E-06 | 0 001480556 | Cd47 | CD47 antigen (Rh-related antigen, integrin-associated signal transducer) |
| ENSMUSG00000042129 | 191 8852336 | -0 477540806 | 0 104747989 | -4 558949635 | 5 14E-06 | 0 001549132 | Rassf4 | Ras association (RalGDS/AF-6) domain family member 4 |
| ENSMUSG00000018707 | 10622 40131 | -0 173071555 | 0 038141474 | -4 537621104 | 5 69E-06 | 0 001675104 | Dync1h1 | dynein cytoplasmic 1 heavy chain 1 |
| ENSMUSG00000027165 | 854 3528383 | -0 257821336 | 0 056835177 | -4 536298673 | 5 73E-06 | 0 001675104 | Iftap | intraflagellar transport associated protein |
| ENSMUSG00000031167 | 6814 431668 | -0 19651911 | 0 043431283 | -4 524828563 | 6 04E-06 | 0 001743305 | Rbm3 | RNA binding motif (RNP1, RRM) protein 3 |
| ENSMUSG00000014633 | 393 7552088 | -0 324777084 | 0 072894068 | -4 455466559 | 8 37E-06 | 0 002380338 | Cmc2 | COX assembly mitochondrial protein 2 |
| ENSMUSG00000056596 | 216 598934 | -0 587592086 | 0 131978637 | -4 452175742 | 8 50E-06 | 0 002383551 | Trnp1 | TMF1-regulated nuclear protein 1 |
| ENSMUSG00000025351 | 157 3330519 | -0 519952292 | 0 11741209 | -4 428439122 | 9 49E-06 | 0 00258958 | Cd63 | CD63 antigen |
| ENSMUSG00000033826 | 2822 845962 | -0 23571229 | 0 053209847 | -4 429862228 | 9 43E-06 | 0 00258958 | Dnah8 | dynein, axonemal, heavy chain 8 |
| ENSMUSG00000024544 | 784 9549093 | 0 260760266 | 0 059383885 | 4 391094773 | 1 13E-05 | 0 003035924 | Ldlrad4 | low density lipoprotein receptor class A domain containing 4 |
| ENSMUSG00000068220 | 1701 954158 | -0 650550579 | 0 148778099 | -4 372623269 | 1 23E-05 | 0 00326112 | Lgals1 | lectin, galactose binding, soluble 1 |
| ENSMUSG00000021457 | 734 6310412 | -0 308847029 | 0 071267443 | -4 333634208 | 1 47E-05 | 0 003845553 | Syk | spleen tyrosine kinase |
| ENSMUSG00000053310 | 1290 74721 | -0 288492517 | 0 066724108 | -4 323662411 | 1 53E-05 | 0 003959639 | Nrgn | neurogranin |
| ENSMUSG00000057738 | 5440 582381 | -0 168144751 | 0 038908495 | -4 321543433 | 1 55E-05 | 0 003959639 | Sptan1 | spectrin alpha, non-erythrocytic 1 |
| ENSMUSG00000071203 | 4514 710548 | -0 390459379 | 0 090423636 | -4 318111893 | 1 57E-05 | 0 00397142 | Naip5 | NLR family, apoptosis inhibitory protein 5 |
| ENSMUSG00000032231 | 3704 814575 | -0 185199835 | 0 042929388 | -4 314057214 | 1 60E-05 | 0 003995065 | Anxa2 | annexin A2 |
| ENSMUSG00000023067 | 31 67440802 | -1 361317443 | 0 317735868 | -4 284431133 | 1 83E-05 | 0 004510699 | Cdkn1a | cyclin-dependent kinase inhibitor 1A (P21) |
| ENSMUSG00000017386 | 420 3036837 | 0 3103847 | 0 072761519 | 4 265780933 | 1 99E-05 | 0 004787777 | Traf4 | TNF receptor associated factor 4 |
| ENSMUSG00000051223 | 7552 551804 | -0 159346812 | 0 037346979 | -4 266658716 | 1 98E-05 | 0 004787777 | Bzw1 | basic leucine zipper and W2 domains 1 |
| ENSMUSG00000001175 | 5515 356307 | -0 203777239 | 0 047911101 | -4 253236395 | 2 11E-05 | 0 005004567 | Calm1 | calmodulin 1 |
| ENSMUSG00000035311 | 1300 887612 | -0 195346771 | 0 046104001 | -4 237089378 | 2 26E-05 | 0 005300935 | Gnptab | N-acetylglucosamine-1-phosphate transferase, alpha and beta subunits |
| ENSMUSG00000089726 | 356 9137159 | -0 348080685 | 0 08218914 | -4 235117716 | 2 28E-05 | 0 005300935 | NA | NA |
| ENSMUSG00000037465 | 331 0813683 | -0 378106278 | 0 08969866 | -4 215294624 | 2 49E-05 | 0 005722946 | Klf10 | Kruppel-like factor 10 |
| ENSMUSG00000020850 | 8865 452626 | -0 139604547 | 0 033199654 | -4 205000097 | 2 61E-05 | 0 00585665 | Prpf8 | pre-mRNA processing factor 8 |
| ENSMUSG00000034220 | 698 2968798 | 0 336959736 | 0 080114683 | 4 205967283 | 2 60E-05 | 0 00585665 | Gpc1 | glypican 1 |
| ENSMUSG00000056413 | 72 2070102 | -0 702309133 | 0 167252827 | -4 199086776 | 2 68E-05 | 0 005881003 | Adap1 | ArfGAP with dual PH domains 1 |
| ENSMUSG00000064065 | 522 3263956 | 0 318677248 | 0 07587 | 4 200306404 | 2 67E-05 | 0 005881003 | Ipcef1 | interaction protein for cytohesin exchange factors 1 |
| ENSMUSG00000103427 | 1614 705311 | -0 292864042 | 0 070117873 | -4 176738825 | 2 96E-05 | 0 00641963 | NA | NA |
| ENSMUSG00000031074 | 54 11055072 | -0 873447306 | 0 210225029 | -4 154820724 | 3 26E-05 | 0 006991895 | Fgf3 | fibroblast growth factor 3 |
| ENSMUSG00000076441 | 586 9514172 | -0 334342939 | 0 08052518 | -4 152029728 | 3 30E-05 | 0 007003233 | Ass1 | argininosuccinate synthetase 1 |
| ENSMUSG00000020901 | 139 2996378 | 0 491106713 | 0 118511325 | 4 143964415 | 3 41E-05 | 0 007178719 | Pik3r5 | phosphoinositide-3-kinase regulatory subunit 5 |
| ENSMUSG00000070639 | 691 9987853 | -0 253694687 | 0 061594516 | -4 118786918 | 3 81E-05 | 0 007913778 | Lrrc8b | leucine rich repeat containing 8 family, member B |
| ENSMUSG00000117485 | 165 7376977 | -0 598640881 | 0 145413615 | -4 116814524 | 3 84E-05 | 0 007913778 | NA | NA |
| ENSMUSG00000065701 | 349 3394562 | -0 403461591 | 0 098090379 | -4 113161698 | 3 90E-05 | 0 007958887 | Rny1 | RNA, Y1 small cytoplasmic, Ro-associated |
| ENSMUSG00000022935 | 50 81629824 | 0 948630537 | 0 230889244 | 4 108595624 | 3 98E-05 | 0 008036683 | Grik1 | glutamate receptor, ionotropic, kainate 1 |
| ENSMUSG00000031431 | 646 7468396 | -0 289697342 | 0 070597823 | -4 103488352 | 4 07E-05 | 0 008134907 | Tsc22d3 | TSC22 domain family, member 3 |
| ENSMUSG00000033256 | 217 3505832 | -0 377088622 | 0 092346955 | -4 083389902 | 4 44E-05 | 0 00878488 | Shf | Src homology 2 domain containing F |
| ENSMUSG00000015619 | 894 7862466 | 0 222758252 | 0 054825885 | 4 063012444 | 4 84E-05 | 0 009482288 | Gata3 | GATA binding protein 3 |
| ENSMUSG00000048126 | 72 80677742 | 0 709714176 | 0 174759974 | 4 061079666 | 4 88E-05 | 0 009482288 | Col6a3 | collagen, type VI, alpha 3 |
| ENSMUSG00000043719 | 23 42665702 | 1 179425354 | 0 291791846 | 4 042009303 | 5 30E-05 | 0 010189702 | Col6a6 | collagen, type VI, alpha 6 |
| ENSMUSG00000009418 | 36 92726337 | 1 159414449 | 0 287961535 | 4 026282358 | 5 67E-05 | 0 010792666 | Nav1 | neuron navigator 1 |
| ENSMUSG00000022372 | 358 5705184 | -0 37123186 | 0 092268463 | -4 023388368 | 5 74E-05 | 0 010824093 | Sla | src-like adaptor |
| ENSMUSG00000054008 | 2164 598875 | -0 161879884 | 0 040324509 | -4 014429147 | 5 96E-05 | 0 011139438 | Ndst1 | N-deacetylase/N-sulfotransferase (heparan glucosaminyl) 1 |
| ENSMUSG00000034765 | 264 9862166 | -0 453790281 | 0 113946757 | -3 982476499 | 6 82E-05 | 0 012632178 | Dusp5 | dual specificity phosphatase 5 |
| ENSMUSG00000020422 | 606 9301928 | 0 276328673 | 0 069454702 | 3 978545222 | 6 93E-05 | 0 012726086 | Tns3 | tensin 3 |
| ENSMUSG00000022508 | 86 57751044 | 0 630100994 | 0 159094178 | 3 960553448 | 7 48E-05 | 0 013600521 | Bcl6 | B cell leukemia/lymphoma 6 |
| ENSMUSG00000030770 | 15 34777855 | 1 607831197 | 0 406310836 | 3 957145746 | 7 59E-05 | 0 013672758 | Parva | parvin, alpha |
| ENSMUSG00000005089 | 25 30349077 | 1 490802078 | 0 378963765 | 3 933890822 | 8 36E-05 | 0 014933028 | Slc1a2 | solute carrier family 1 (glial high affinity glutamate transporter), member 2 |
| ENSMUSG00000030209 | 40 11834233 | 0 969045423 | 0 24717117 | 3 9205439 | 8 83E-05 | 0 015646358 | Grin2b | glutamate receptor, ionotropic, NMDA2B (epsilon 2) |
| ENSMUSG00000037337 | 1047 434357 | -0 221391389 | 0 056505842 | -3 918026512 | 8 93E-05 | 0 01567314 | Map4k1 | mitogen-activated protein kinase kinase kinase kinase 1 |
| ENSMUSG00000020638 | 211 3394396 | -0 509639443 | 0 130236156 | -3 913194757 | 9 11E-05 | 0 015852371 | Cmpk2 | cytidine monophosphate (UMP-CMP) kinase 2, mitochondrial |
| ENSMUSG00000045312 | 23 26053874 | 1 267648889 | 0 324212227 | 3 909935476 | 9 23E-05 | 0 015909329 | Lhfpl2 | lipoma HMGIC fusion partner-like 2 |
| ENSMUSG00000061462 | 65 8810231 | 0 7530033 | 0 192672663 | 3 908199987 | 9 30E-05 | 0 015909329 | Obscn | obscurin, cytoskeletal calmodulin and titin-interacting RhoGEF |
| ENSMUSG00000025586 | 490 4341103 | -0 281281437 | 0 072189319 | -3 896441202 | 9 76E-05 | 0 016423161 | Cpeb1 | cytoplasmic polyadenylation element binding protein 1 |

| | | | | | | | | |
|---|---|---|---|---|---|---|---|---|
| ENSMUSG00000035064 | 1934 800434 | -0 204446699 | 0 052469988 | -3 896450299 | 9 76E-05 | 0 016423161 | Eef2k | eukaryotic elongation factor-2 kinase |
| ENSMUSG00000063953 | 585 9141858 | -0 240480418 | 0 06182658 | -3 889595968 | 0 000100411 | 0 016753742 | Amd2 | S-adenosylmethionine decarboxylase 2 |
| ENSMUSG00000034041 | 348 1057181 | -0 318721135 | 0 08202621 | -3 885601123 | 0 000102077 | 0 016875743 | Lyl1 | lymphoblastomic leukemia 1 |
| ENSMUSG00000067919 | 23 38875167 | 1 079912416 | 0 278051849 | 3 883852665 | 0 000102814 | 0 016875743 | Rex2 | reduced expression 2 |
| ENSMUSG00000026547 | 2884 533758 | -0 178628269 | 0 046341818 | -3 8545805 | 0 000115928 | 0 0188748 | Tagln2 | transgelin 2 |
| ENSMUSG00000032053 | 1186 686676 | -0 195337352 | 0 050704584 | -3 852459403 | 0 000116937 | 0 018886788 | Pou2af1 | POU domain, class 2, associating factor 1 |
| ENSMUSG00000030043 | 11 27160915 | 1 836723845 | 0 477413661 | 3 84723772 | 0 000119457 | 0 019138904 | Tacr1 | tachykinin receptor 1 |
| ENSMUSG00000033335 | 1803 362255 | -0 192692523 | 0 050110896 | -3 845321836 | 0 000120394 | 0 019138904 | Dnm2 | dynamin 2 |
| ENSMUSG00000028788 | 2911 127626 | -0 15648993 | 0 040729216 | -3 842203384 | 0 000121935 | 0 019140048 | Ptp4a2 | protein tyrosine phosphatase 4a2 |
| ENSMUSG00000038146 | 1828 122737 | -0 242565465 | 0 063143849 | -3 841474195 | 0 000122298 | 0 019140048 | Notch3 | notch 3 |
| ENSMUSG00000020844 | 556 5960603 | -0 278480757 | 0 072740076 | -3 828436429 | 0 00012896 | 0 01957573 | Nxn | nucleoredoxin |
| ENSMUSG00000021500 | 4347 325027 | -0 149607455 | 0 039061072 | -3 830090882 | 0 000128096 | 0 01957573 | Ddx46 | DEAD box helicase 46 |
| ENSMUSG00000026305 | 1949 639366 | -0 173777633 | 0 045376233 | -3 829705995 | 0 000128296 | 0 01957573 | Lrrfip1 | leucine rich repeat (in FLII) interacting protein 1 |
| ENSMUSG00000035561 | 2166 666069 | -0 200959477 | 0 052472091 | -3 829835522 | 0 000128229 | 0 01957573 | Aldh1b1 | aldehyde dehydrogenase 1 family, member B1 |
| ENSMUSG00000064341 | 14778 42147 | -0 205812629 | 0 053868881 | -3 820621909 | 0 000133116 | 0 020055751 | ND1 | NADH dehydrogenase subunit 1 |
| ENSMUSG00000075232 | 1123 057656 | -0 193089975 | 0 050738744 | -3 805572626 | 0 000141476 | 0 021157547 | Amd1 | S-adenosylmethionine decarboxylase 1 |
| ENSMUSG00000020363 | 11 17728156 | 1 554563227 | 0 409363206 | 3 797515763 | 0 000146153 | 0 021380971 | Gfpt2 | glutamine fructose-6-phosphate transaminase 2 |
| ENSMUSG00000041482 | 31 33838552 | 1 173456306 | 0 308817966 | 3 799831732 | 0 000144794 | 0 021380971 | Piezo2 | piezo-type mechanosensitive ion channel component 2 |
| ENSMUSG00000066007 | 26 7348062 | 1 039478971 | 0 273854459 | 3 795735054 | 0 000147207 | 0 021380971 | Zfp600 | zinc finger protein 600 |
| ENSMUSG00000104988 | 10 35424739 | 1 969422853 | 0 518541216 | 3 79800639 | 0 000145865 | 0 021380971 | NA | NA |
| ENSMUSG00000070733 | 4680 631757 | -0 156135808 | 0 041181074 | -3 791445789 | 0 000149773 | 0 021598314 | Fryl | FRY like transcription coactivator |
| ENSMUSG00000020745 | 1814 471137 | -0 177370093 | 0 046876417 | -3 783780978 | 0 000154464 | 0 021668187 | Pafah1b1 | platelet-activating factor acetylhydrolase, isoform 1b, subunit 1 |
| ENSMUSG00000028736 | 81 74727738 | 0 641698956 | 0 169598248 | 3 783641417 | 0 00015455 | 0 021668187 | Pax7 | paired box 7 |
| ENSMUSG00000035891 | 104 1383453 | 0 559654994 | 0 147826687 | 3 785886077 | 0 000153162 | 0 021668187 | Cerk | ceramide kinase |
| ENSMUSG00000118590 | 28 88450069 | 1 107570433 | 0 292469367 | 3 786962189 | 0 0001525 | 0 021668187 | NA | NA |
| ENSMUSG00000015839 | 890 9036854 | -0 198897921 | 0 052706533 | -3 773686275 | 0 000160853 | 0 022067782 | Nfe2l2 | nuclear factor, erythroid derived 2, like 2 |
| ENSMUSG00000025203 | 6578 292181 | 0 15539237 | 0 04119948 | 3 771707028 | 0 000162135 | 0 022067782 | Scd2 | stearoyl-Coenzyme A desaturase 2 |
| ENSMUSG00000028991 | 4878 06508 | -0 163365237 | 0 043340179 | -3 769371505 | 0 000163659 | 0 022067782 | Mtor | mechanistic target of rapamycin kinase |
| ENSMUSG00000030536 | 6388 05877 | -0 141168411 | 0 037455985 | -3 768914618 | 0 000163959 | 0 022067782 | Iqgap1 | IQ motif containing GTPase activating protein 1 |
| ENSMUSG00000063531 | 10 36684582 | 1 719294303 | 0 455204653 | 3 776969968 | 0 000158748 | 0 022067782 | Sema3e | sema domain, immunoglobulin domain (Ig), short basic domain, secreted, (semaphorin) 3E |
| ENSMUSG00000104145 | 812 6431926 | -0 299467083 | 0 079404468 | -3 771413514 | 0 000162325 | 0 022067782 | NA | NA |
| ENSMUSG00000009376 | 10 04421947 | 1 881061574 | 0 499620451 | 3 764981139 | 0 000166562 | 0 022269624 | Met | met proto-oncogene |
| ENSMUSG00000022449 | 195 3478912 | 0 391169872 | 0 104050964 | 3 759406508 | 0 000170317 | 0 022328104 | Adamts20 | a disintegrin-like and metallopeptidase (reprolysin type) with thrombospondin type 1 motif, 20 |
| ENSMUSG00000025810 | 1178 651148 | -0 290803398 | 0 077301774 | -3 761923995 | 0 000168611 | 0 022328104 | Nrp1 | neuropilin 1 |
| ENSMUSG00000068015 | 851 4134692 | -0 228642465 | 0 060805914 | -3 760201082 | 0 000169777 | 0 022328104 | Lrch1 | leucine-rich repeats and calponin homology (CH) domain containing 1 |
| ENSMUSG00000048148 | 1057 416039 | -0 267960456 | 0 071326076 | -3 756837222 | 0 000172074 | 0 022412961 | Nwd1 | NACHT and WD repeat domain containing 1 |
| ENSMUSG00000026024 | 1456 939609 | -0 202841515 | 0 054062987 | -3 75194799 | 0 000175466 | 0 022708208 | Als2 | alsin Rho guanine nucleotide exchange factor |
| ENSMUSG00000061751 | 41 68110646 | 1 034265911 | 0 27596658 | 3 747794069 | 0 000178397 | 0 022940442 | Kalrn | kalirin, RhoGEF kinase |
| ENSMUSG00000022132 | 141 5848434 | -0 575622369 | 0 153944152 | -3 739163588 | 0 000184634 | 0 023443816 | Cldn10 | claudin 10 |
| ENSMUSG00000032816 | 22 33006921 | 1 241905784 | 0 331998305 | 3 740699176 | 0 000183509 | 0 023443816 | Igdcc4 | immunoglobulin superfamily, DCC subclass, member 4 |
| ENSMUSG00000047904 | 98 00708848 | -0 571030851 | 0 153414255 | -3 722149888 | 0 000197534 | 0 024915546 | Sstr2 | somatostatin receptor 2 |
| ENSMUSG00000096878 | 50 29275562 | 0 983512874 | 0 264337352 | 3 720673094 | 0 000198693 | 0 024915546 | Gm21083 | predicted gene, 21083 |
| ENSMUSG00000003660 | 5463 963053 | -0 140143862 | 0 037717913 | -3 715578333 | 0 000202739 | 0 02526609 | Snrnp200 | small nuclear ribonucleoprotein 200 (U5) |
| ENSMUSG00000061601 | 28 61430733 | 1 317531961 | 0 355328131 | 3 707930345 | 0 00020896 | 0 025881569 | Pclo | piccolo (presynaptic cytomatrix protein) |
| ENSMUSG00000069601 | 61 60421914 | 0 890273389 | 0 240297666 | 3 704877381 | 0 000211493 | 0 026035564 | Ank3 | ankyrin 3, epithelial |
| ENSMUSG00000108095 | 44 49753787 | 0 76126895 | 0 205728864 | 3 70035072 | 0 000215302 | 0 026343794 | NA | NA |
| ENSMUSG00000058013 | 6047 527188 | -0 128946556 | 0 035010059 | -3 683128766 | 0 000230389 | 0 027852201 | Septin11 | septin 11 |
| ENSMUSG00000068587 | 24 45991167 | 1 13687528 | 0 308652004 | 3 683356228 | 0 000230183 | 0 027852201 | Mgam | maltase-glucoamylase |
| ENSMUSG00000038280 | 545 4590272 | -0 239054487 | 0 06493231 | -3 681595276 | 0 000231779 | 0 027853507 | Ostm1 | osteopetrosis associated transmembrane protein 1 |
| ENSMUSG00000007817 | 4608 929229 | -0 147297326 | 0 040089894 | -3 674175986 | 0 000238618 | 0 028505701 | Zmiz1 | zinc finger, MIZ-type containing 1 |
| ENSMUSG00000026832 | 734 7034379 | 0 216548375 | 0 059000115 | 3 670304254 | 0 000242262 | 0 028770742 | Cytip | cytohesin 1 interacting protein |
| ENSMUSG00000032601 | 1226 105284 | -0 194395109 | 0 053188266 | -3 654849495 | 0 000257333 | 0 030381835 | Prkar2a | protein kinase, cAMP dependent regulatory, type II alpha |
| ENSMUSG00000013089 | 170 7572102 | 0 441705491 | 0 12116969 | 3 645346393 | 0 000267032 | 0 03088526 | Etv5 | ets variant 5 |
| ENSMUSG00000040219 | 572 4153793 | -0 271827642 | 0 074582352 | -3 644664398 | 0 000267741 | 0 03088526 | Ttc12 | tetratricopeptide repeat domain 12 |
| ENSMUSG00000072949 | 65 21735681 | -0 63424529 | 0 173995384 | -3 645184574 | 0 0002672 | 0 03088526 | Acot1 | acyl-CoA thioesterase 1 |
| ENSMUSG00000098145 | 65 71992667 | -0 568272593 | 0 155794583 | -3 647576066 | 0 000264726 | 0 03088526 | NA | NA |
| ENSMUSG00000103187 | 21 07345479 | 1 167238044 | 0 320386155 | 3 64322249 | 0 000269246 | 0 03088526 | NA | NA |
| ENSMUSG00000043535 | 4321 539255 | -0 185565048 | 0 051074804 | -3 633201361 | 0 000279926 | 0 031929006 | Setx | senataxin |
| ENSMUSG00000039989 | 1485 378983 | -0 26536348 | 0 073082875 | -3 630993972 | 0 000282332 | 0 032022451 | Cbx4 | chromobox 4 |
| ENSMUSG00000045382 | 1351 445782 | -0 191144515 | 0 052712803 | -3 626149681 | 0 000287679 | 0 032446615 | Cxcr4 | chemokine (C-X-C motif) receptor 4 |
| ENSMUSG00000023132 | 1116 50323 | -0 253644935 | 0 070119497 | -3 617323963 | 0 000297665 | 0 033386397 | Gzma | granzyme A |
| ENSMUSG00000008855 | 312 5493578 | 0 339798372 | 0 094014216 | 3 614329688 | 0 000301126 | 0 033587991 | Hdac5 | histone deacetylase 5 |
| ENSMUSG00000026825 | 18 29456816 | 1 260834805 | 0 349310804 | 3 609492719 | 0 000306796 | 0 034032485 | Dnm1 | dynamin 1 |
| ENSMUSG00000034807 | 1101 769552 | -0 238499869 | 0 066110259 | -3 607607541 | 0 000309033 | 0 034093315 | Colgalt1 | collagen beta(1-O)galactosyltransferase 1 |
| ENSMUSG00000042793 | 14 10451338 | 1 472693948 | 0 409106851 | 3 599778261 | 0 000318489 | 0 034945475 | Lgr6 | leucine-rich repeat-containing G protein-coupled receptor 6 |
| ENSMUSG00000114576 | 61 14605007 | -0 659258642 | 0 183233705 | -3 59791144 | 0 000320783 | 0 03500694 | NA | NA |
| ENSMUSG00000021313 | 58 18635872 | 0 71616079 | 0 199496625 | 3 58983913 | 0 000330882 | 0 035914939 | Ryr2 | ryanodine receptor 2, cardiac |
| ENSMUSG00000042744 | 3910 224586 | -0 154869413 | 0 043270811 | -3 579073523 | 0 000344814 | 0 037135479 | Hectd4 | HECT domain E3 ubiquitin protein ligase 4 |
| ENSMUSG00000079429 | 336 3976951 | 0 294099885 | 0 082189301 | 3 578323226 | 0 000345806 | 0 037135479 | Mroh2a | maestro heat-like repeat family member 2A |
| ENSMUSG00000034156 | 758 2100031 | 0 209542672 | 0 058670597 | 3 571510818 | 0 000354928 | 0 037913434 | Tspoap1 | TSPO associated protein 1 |
| ENSMUSG00000110618 | 101 3682684 | -0 54114341 | 0 151652715 | -3 568306766 | 0 000359296 | 0 038178004 | Gm39822 | predicted gene, 39822 |
| ENSMUSG00000091255 | 31 7434741 | 1 099864327 | 0 30937863 | 3 555075305 | 0 000377871 | 0 039941542 | Speer4e | spermatogenesis associated glutamate (E)-rich protein 4e |
| ENSMUSG00000033653 | 610 4074994 | -0 220461698 | 0 06204258 | -3 553393427 | 0 000380295 | 0 039988442 | Vps8 | VPS8 CORVET complex subunit |
| ENSMUSG00000041828 | 8 743768143 | 1 690875659 | 0 476677191 | 3 547213274 | 0 000389329 | 0 040726263 | Abca8a | ATP-binding cassette, sub-family A (ABC1), member 8a |
| ENSMUSG00000052469 | 18 76106858 | 1 275312731 | 0 360122852 | 3 541326866 | 0 00039812 | 0 041431163 | Tcp10c | t-complex protein 10c |
| ENSMUSG00000019891 | 96 95890985 | 0 519869188 | 0 147475055 | 3 525133044 | 0 00042327 | 0 041701169 | Dcbld1 | discoidin, CUB and LCCL domain containing 1 |
| ENSMUSG00000024360 | 2520 727413 | -0 168427278 | 0 047698185 | -3 531104531 | 0 000413828 | 0 041701169 | Etf1 | eukaryotic translation termination factor 1 |
| ENSMUSG00000028528 | 1713 613868 | -0 172673441 | 0 048933254 | -3 528754536 | 0 00041752 | 0 041701169 | Dnajc6 | DnaJ heat shock protein family (Hsp40) member C6 |
| ENSMUSG00000030213 | 3218 558472 | -0 137481016 | 0 038930356 | -3 531460507 | 0 000413272 | 0 041701169 | Atf7ip | activating transcription factor 7 interacting protein |
| ENSMUSG00000030589 | 132 2230628 | -0 483731833 | 0 136922396 | -3 532890516 | 0 000411043 | 0 041701169 | Rasgrp4 | RAS guanyl releasing protein 4 |
| ENSMUSG00000041870 | 5517 523677 | -0 166336908 | 0 047186063 | -3 525127913 | 0 000423278 | 0 041701169 | Ankrd13a | ankyrin repeat domain 13a |
| ENSMUSG00000046634 | 21 42495018 | 1 124582309 | 0 31868528 | 3 52881787 | 0 00041742 | 0 041701169 | Pkd1l1 | polycystic kidney disease 1 like 1 |
| ENSMUSG00000052105 | 16 21372026 | 1 210166655 | 0 342682195 | 3 531454726 | 0 000413281 | 0 041701169 | Mtcl1 | microtubule crosslinking factor 1 |
| ENSMUSG00000054036 | 12 18408724 | 1 408884602 | 0 399680222 | 3 525029577 | 0 000423436 | 0 041701169 | Olfr279 | olfactory receptor 279 |
| ENSMUSG00000060096 | 202 5210973 | -0 353507367 | 0 099988876 | -3 535466942 | 0 000407055 | 0 041701169 | NA | NA |
| ENSMUSG00000062380 | 339 3673222 | -0 333946783 | 0 094425213 | -3 536627259 | 0 000405271 | 0 041701169 | Tubb3 | tubulin, beta 3 class III |
| ENSMUSG00000026116 | 2238 513478 | -0 240329114 | 0 068244377 | -3 521595834 | 0 000428958 | 0 041824465 | Tmem131 | transmembrane protein 131 |
| ENSMUSG00000034282 | 545 4353341 | -0 270127393 | 0 076759552 | -3 51913718 | 0 000432953 | 0 041824465 | Evpl | envoplakin |
| ENSMUSG00000054263 | 15 25487244 | 1 485698644 | 0 422178526 | 3 519124143 | 0 000432974 | 0 041824465 | Lifr | LIF receptor alpha |
| ENSMUSG00000112522 | 13 15277562 | 1 356701854 | 0 385236564 | 3 521736978 | 0 000428729 | 0 041824465 | NA | NA |
| ENSMUSG00000059854 | 44 25269797 | 0 92439981 | 0 26289017 | 3 516296598 | 0 000437612 | 0 042071167 | Hydin | HYDIN, axonemal central pair apparatus protein |
| ENSMUSG00000034361 | 260 9171454 | -0 420877581 | 0 119736991 | -3 515017174 | 0 000439726 | 0 042074024 | Cpne2 | copine II |
| ENSMUSG00000034774 | 68 8946702 | 0 668866136 | 0 190435586 | 3 512295936 | 0 000444253 | 0 042306718 | Dsg1c | desmoglein 1 gamma |
| ENSMUSG00000037458 | 1421 69282 | -0 166739801 | 0 047512172 | -3 509412315 | 0 000449098 | 0 042567332 | Azin1 | antizyme inhibitor 1 |
| ENSMUSG00000052374 | 289 7929459 | 0 323208069 | 0 092141479 | 3 507736947 | 0 000451936 | 0 04263612 | Actn2 | actinin alpha 2 |
| ENSMUSG00000021198 | 23 70187431 | 1 072362989 | 0 306002923 | 3 50442074 | 0 000457602 | 0 042770935 | Unc79 | unc-79 homolog |
| ENSMUSG00000039115 | 4817 561588 | -0 195139638 | 0 055673015 | -3 505102753 | 0 000456431 | 0 042770935 | Itga9 | integrin alpha 9 |
| ENSMUSG00000103385 | 20 81823166 | 1 126193794 | 0 322107889 | 3 496324782 | 0 000471714 | 0 043886809 | NA | NA |
| ENSMUSG00000020340 | 4250 512845 | -0 145686539 | 0 041738987 | -3 490418682 | 0 000482264 | 0 044526489 | Cyfip2 | cytoplasmic FMR1 interacting protein 2 |
| ENSMUSG00000083282 | 33 04515817 | 0 824130376 | 0 236139766 | 3 490010985 | 0 000483001 | 0 044526489 | Ctsf | cathepsin F |
| ENSMUSG00000029761 | 31 48018141 | 0 979425998 | 0 280852114 | 3 48733711 | 0 000487856 | 0 044711536 | Cald1 | caldesmon 1 |
| ENSMUSG00000034951 | 332 0657886 | -0 265964273 | 0 076284651 | -3 486471631 | 0 000489437 | 0 044711536 | Cog7 | component of oligomeric golgi complex 7 |
| ENSMUSG00000070933 | 47 58028931 | 0 963437258 | 0 276865363 | 3 479804222 | 0 00050178 | 0 045632625 | Speer4d | spermatogenesis associated glutamate (E)-rich protein 4D |
| ENSMUSG00000047746 | 19 25695375 | 1 113200171 | 0 320205767 | 3 476515061 | 0 000507976 | 0 045988885 | Fbxo40 | F-box protein 40 |
| ENSMUSG00000030577 | 14 13825474 | 1 430381097 | 0 413007785 | 3 463327206 | 0 000533539 | 0 048087598 | Cd22 | CD22 antigen |
| ENSMUSG00000021556 | 31 48151734 | 0 964912204 | 0 279128696 | 3 456872115 | 0 000546484 | 0 048603383 | Golm1 | golgi membrane protein 1 |
| ENSMUSG00000022472 | 4463 562972 | -0 157462514 | 0 045539343 | -3 457724797 | 0 000544758 | 0 048603383 | Desi1 | desumoylating isopeptidase 1 |
| ENSMUSG00000063684 | 205 5307202 | -0 328812599 | 0 095116172 | -3 456957857 | 0 00054631 | 0 048603383 | NA | NA |

**Supplemental Table 5. Characteristics of patient samples.**

| identifier | use | type | age at diagnos | sex | cytogenetic | immunophenotype | timing of sample |
|---|---|---|---|---|---|---|---|
| P2 | Xenograft transplantation | bone marrow | 1 | F | t(4;11) | B cell ALL | diagnosis |
| P6 | Xenograft transplantation | bone marrow | 1 | F | t(4;11) | B cell ALL | diagnosis |
| P10 | Xenograft transplantation | bone marrow | 1 | F | t(4;11) | B cell ALL | relapse |
| 0868 | miR validation | bone marrow + cells from CSF | 1 | M | t(11q23) | B cell ALL | diagnosis |
| 1057 | miR validation | bone marrow + cells from CSF | 2 | M | t(11q23) | B cell ALL | diagnosis |
| 1615 | miR validation | bone marrow + cells from CSF | 7 | F | t(4;11) | B cell ALL | day 35 induction |
| 1737 | miR validation | bone marrow + cells from CSF | 2 | F | t(11q23) | B cell ALL | day 30 induction |
| 1741 | miR validation | cells from CSF | 1 | M | t(11q23) | B cell ALL | diagnosis |
| 1889 | miR validation | bone marrow + cells from CSF | 7 | F | t(11q23) | B cell ALL | diagnosis |
